# Transcriptional response to Phytophthora root rot in raspberry identifies *RiABP19*, a Germin-like protein (GLP) gene with a putative role in resistance

**DOI:** 10.1101/2025.07.01.662284

**Authors:** Raisa Osama, Charlene E. Ogilvie, S. Ronan Fisher, Lydia Welsh, John Fuller, Kay Smith, Aurélia Bézanger, Linda Milne, Hazel McLellan, Craig G. Simpson, Eleanor M. Gilroy, Murray R. Grant

## Abstract

Most phytophthora root rot (PRR) outbreaks in symptomatic commercially cultivated raspberry varieties are associated with the prevalence of *Phytophthora rubi*. Reduced availability of chemical actives and the persistent presence of *Phytophthora* oospores in the soil contribute to its devastating impact on raspberry-growing regions. In this study, we examined the variation in root morphology in two contrasting raspberry cultivars, Latham (PRR resistant) and Glen Moy (PRR susceptible). We performed RNA-sequencing on Latham roots challenged with *P. rubi,* to study the transcriptomic response and uncover mechanisms underpinning resistance. We established a new raspberry reference transcript dataset that allowed quantification of raspberry root gene expression. Transcripts significantly upregulated in Latham challenged with *P. rubi*, included many with characterised roles in resistance, such as Pathogenesis-related proteins and a Germin-like protein, designated *RiABP19.* The homologous Glen Moy *RiABP19* gene showed no differential transcriptional response to PRR infection, indicating a resistance cultivar-specific induction signature following PRR challenge. Three-dimensional structural modelling predicts that RiABP19 contains conserved active sites implicated in auxin-binding and superoxide dismutase activity and can form a homo-hexamer like true germins. Co-immunoprecipitation assays confirmed that RiABP19 can form both homo- and heterodimers *in planta*. Virus-induced gene silencing of *RiABP19* orthologs of in the model plant *Nicotiana benthamiana* strongly impacts immune signalling, enhancing *Phytophthora infestans* colonization and attenuating resistance and cell death triggered by the tomato Cf4/Avr4 interaction. These findings suggest that RiABP19 functions as a positive regulator of immunity and may represent a target for future crop improvement in raspberries.

## Introduction

Field production of red raspberries (*Rubus idaeus)* is severely threatened by PRR, as large areas of land used for soft fruit growing are contaminated by *P. rubi* oospores. These thick-walled oospores can persist in soil for over a decade (Newton et al., 2010). In the presence of susceptible raspberry cultivars and increasingly wet summers, growers have rapidly adopted pot-based growing systems, to limit waterlogging and contact with contaminated soil (Hoashi-Erhardt et al 2008). Currently, PRR can be controlled by prophylactic chemicals, but the limited window for application, diminishing number of actives approved for use on growing crops and reports of loss of sensitivity to actives mean growers need more commercially viable PRR-resistant raspberry cultivars.

Latham is a hardy cultivar of red raspberry derived from European wild red raspberry *Rubus idaeus* and the American wild red raspberry *R. strigosus,* bred at the University of Minnesota in 1918 (Jennings 2018). Latham roots initially exhibit signs of infection upon exposure to *P. rubi*, however, their root vigour and resistance to PRR enable plants to overcome the disease (Laun and Zinkernagel, 1997). Accordingly, Latham has been utilised as a genetic source for PRR resistance in breeding programmes and as a tool to study molecular mechanisms behind its underlying resistance (Graham et al., 2011). Using a segregating population, derived from a cross between PRR-resistant cv. Latham and PRR-susceptible cv. Glen Moy, identified two specific QTLs associated with PRR resistance in Latham (Graham et al., 2011). One of the QTL markers, on linkage group 6, was associated with genes involved in root development, stress response and defence, including a gene resembling a germin-like or auxin-binding gene (Graham et al., 2011). Subsequent development of a simple sequence repeat marker (Rub118b) that distinguishes resistant and susceptible germplasm has since been instrumental in raspberry breeding programmes, leading to release of a PRR-resistant cultivar, Glen Mor. However, the molecular mechanisms underpinning resistance in Latham remains to be elucidated.

Auxins are tightly regulated growth-promoting phytohormones that initiate root growth and development via establishment of localised auxin gradients (Blakeslee et al., 2019). Misregulation or inhibition of either auxin synthesis or transport results in significant disruptions to root or shoot apical meristem identity, stem or shoot growth, and lateral root or root hair growth (Shigenaga et al., 2016). Auxins are also involved in crosstalk between growth, development and responses to both biotic and abiotic stresses (reviewed in Jing et al., 2023; Kunkel and Johnson, 2021). For example, phytohormones salicylic acid (SA) and jasmonic acid (JA) act as key modulators for responses against biotrophic and necrotropic pathogens respectively and downregulate growth processes via repression of numerous genes involved in auxin signalling (Huot et al., 2014). Conversely, upregulation of auxin signalling has been shown to increase susceptibility to infection, with several pathogens and beneficial microbes found to synthesise auxin to promote interactions with the plant (Kidd et al., 2011). Ultimately timing, location and concentration of auxin accumulation in plant tissues influence the outcome of plant-pathogen interactions by impacting defence signalling on multiple levels (Gilroy and Breen, 2022).

Distinct from auxin transporters, auxin-binding proteins (ABPs) function in auxin receptor systems. AtABP1 predominantly localizes to the endoplasmic reticulum (ER) and cell surface, influencing auxin-dependent cell expansion and division (Grones and Friml, 2015). ABPs belong to a large family of germin-like proteins (GLPs), conserved in all plants (Dunwell et al., 2008). True germins (GERs) are a group of homologous proteins only found in cereals, originally isolated from germinating wheat embryos and characterised as homohexamer glycoproteins (Thompson and Lane, 1980; Woo et al., 2000). Both GERs and GLPs belong to a biochemically diverse superfamily of proteins called cupins, which are glycoproteins, characterised by the presence of a conserved β-barrel core involved in manganese ion binding (Barman and Banerjee, 2015). Many GERs produce hydrogen peroxide (H_2_O_2_) through oxalate oxidase (OXO) activity and are involved in defence reactions via production of active oxygen species for cross-linking reactions (Otte and Barz, 1996) and stress-related signalling by enhancing extracellular Ca^2+^ levels (Bayles and Aist, 1987). Other germin family members have been shown to possess superoxide dismutase (SOD) activity, such as HvGER5, which is an extra-cellular SOD with antifungal activity against powdery mildew (Zimmermann et al., 2006). Transgenic plants overexpressing GERs and GLPs with OXO/SOD activity have been shown to increase host resistance against different biotic and abiotic stresses (Ilyas et al., 2016). Importantly, OXOs and oxidase-like proteins are classified as Pathogenesis-Related (PR) Proteins, PR-15 and PR-16 respectively (Zribi et al., 2021).

We used RNA sequencing (RNA-seq) to investigate molecular mechanisms underpinning PRR resistance in the roots of cv. Latham after challenge with *P. rubi*. Transcriptional responses in roots at 7 dpi revealed a range of significantly induced genes with putative roles in host defences in addition to genes associated with responses to auxin. Of particular interest, was an *auxin-binding protein 19* (*RiABP19*) ortholog, with a putative role in both defence and responses to auxin that had been previously shown to be associated with a resistance QTL in raspberry (Graham et al., 2011). Genomic analysis of Latham identified multiple *RiABP19* orthologs, but only one showed significant upregulation after challenge with *P. rubi*. We analysed the predicted protein structure of this induced *RiABP19* ortholog to understand its putative function and used co-immunoprecipitation (Co-IP) to confirm that RiABP19 can form both homo- and heterodimers *in planta*. We identified and silenced tobacco ABP19 orthologs in *Nicotiana benthamiana* and found compromised host resistance and increased susceptibility to *P. infestans* infection. This highlights RiABP19 as a positive regulator in host resistance responses to *Phytophthora* pathogens.

## Results

### Role of root morphology of PRR-resistant and susceptible cultivars of raspberry

The root morphology of raspberry cultivars cv. Latham (PRR resistant) and cv. G. Moy (PRR susceptible) are distinct. Latham exhibits a robust root system with numerous primary, secondary and tertiary lateral roots, largely arising from the root-hypocotyl junction. Conversely, G. Moy has a substantial primary root with a weak, unbranched lateral root system and more symptomatic roots darken with necrotic rot after infection **(Fig. 1A)**. Latham displayed an abundance of healthy root growth even at 28 weeks after challenge with *P. rubi* in pot-based, substrate infection assays **(Fig. 1B)**. A cross-section analysis of the primary roots showed roots of Latham featured a thicker cortex with a well-defined stele and abundant root hairs, contrasting with the root structure of G. Moy, which lacks root hairs and has a less well-defined stele **(Fig. 1C)**. These differences in root morphology may contribute to the resistant phenotype of Latham, particularly in hypoxic conditions during our hydroponic infection assays.

**Figure 1:**
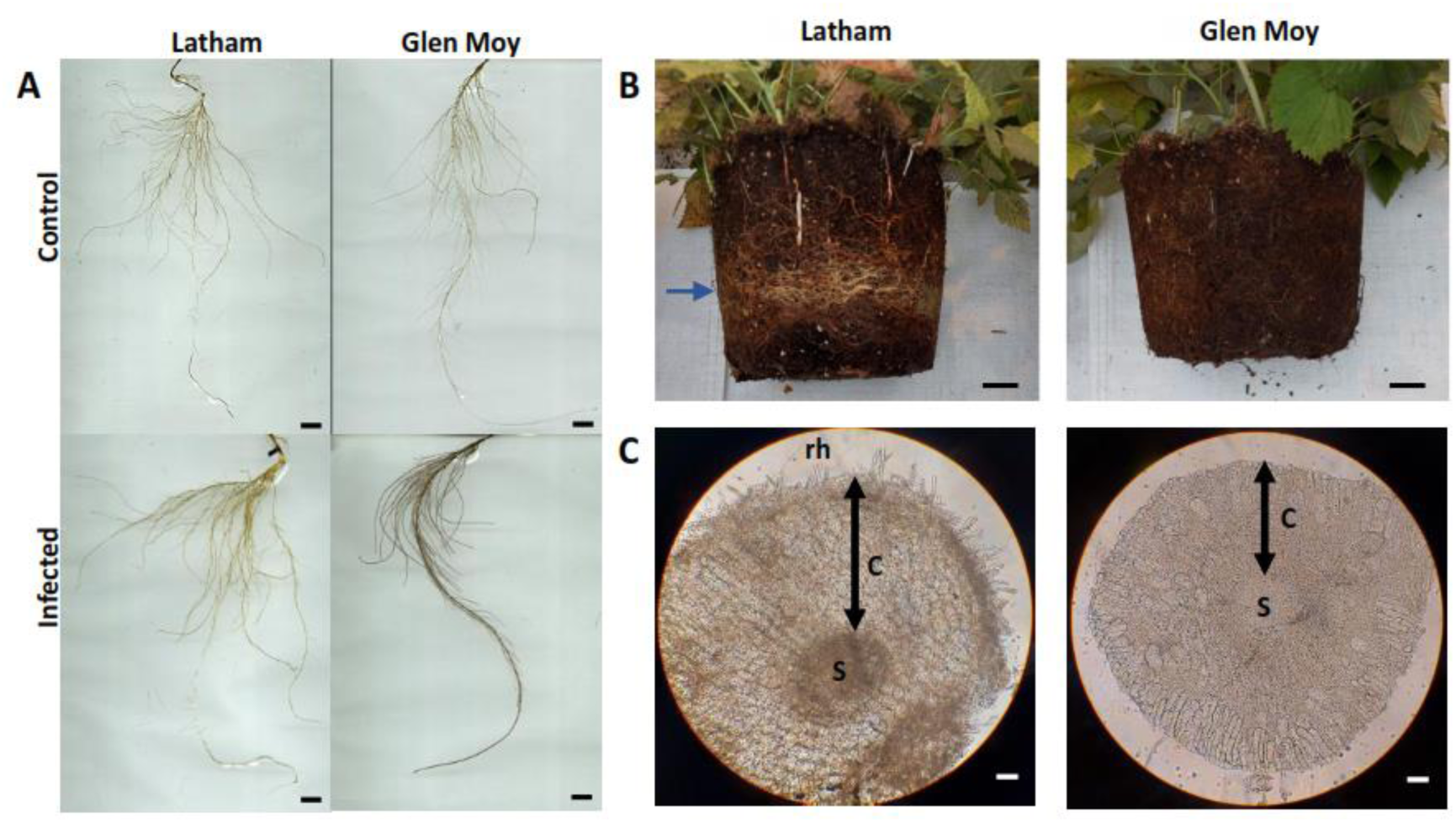
Root morphology and phenotypes of PRR-resistant Latham and susceptible Glen Moy. **A.** Control and infected roots of cv. Latham and cv. G. Moy grown in hydroponics highlight differences in root architecture and signs of root rot at 7 days post-infection. G. Moy displays characteristic darkening necrosis of root rot. Scale bar is equal to 1 cm. **B.** Root balls of infected plants in soil substrate. Growth of new, pale-coloured roots can be seen in the root ball of Latham, as indicated by the blue arrow at 28 weeks post-infection. Scale bar is equal to 5 cm. **C.** Cross sections of the primary root in Latham and G. Moy observed under the light microscope at 40 X magnification. Double-ended arrows indicate thickness of cortex. Root segments were taken from 1.5-2 cm above the tip. **S**: stele; **C**: cortex and **rh**: root hair. Scale bar is equal to 100 μm.

### RNA sequencing (RNA-seq) analysis of Latham roots challenged with *P. rubi*

To uncover evidence of molecular mechanisms underpinning observed PRR resistance in Latham, we performed RNA-seq to study differential expression of genes (DEGs) in raspberry root tips challenged with *P. rubi* at 7 dpi. To do this we first created a raspberry reference transcript dataset (RiRTD) from publicly available and in-house raspberry RNA-sequencing datasets, including the root tip infected RNA-sequencing dataset produced here (see Methods for assembly of RiRTD4). The final RiRTD4 contained 38,976 genes and 141,513 transcript models and was used to identify significant DEGs (see Methods). Overall analysis of the variation between the control and infected Latham samples shows significant differences between infected and uninfected root samples **(Fig. 2A)**. Of 20,463 expressed genes, 1178 were significant DEGs of which, 456 and 722 were upregulated and downregulated, respectively **(Fig. 2B).** GO (Gene Ontology) analysis was used to analyse functional categorisations of DEGs **(Fig. 2C)**. The largest functional category of DEG associated with cellular components was ‘integral component of membrane’ which comprised 618 (46%) of all DEGs, indicating extensive modification of membrane-associated processes in Latham roots as part of defence reprograming induced by *P. rubi*. Based on gene count numbers, the top functional biological process categories were ‘protein phosphorylation’ and ‘defence response’. In ‘defence response’, most genes upregulated are *PR* genes, supporting induction of an active immune response in Latham to infection **(Supplementary Table S1)**.

**Figure 2:**
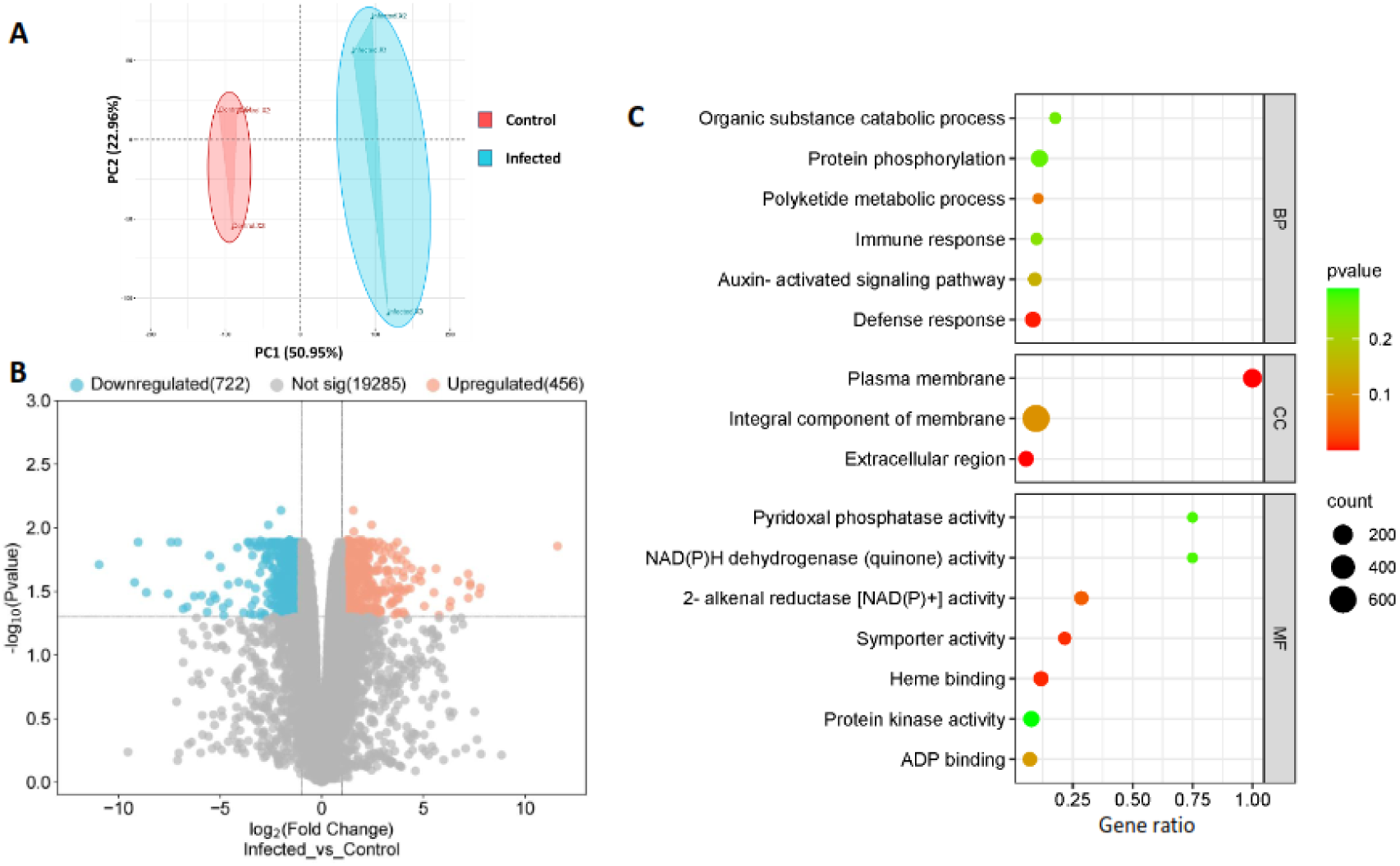
Global Transcriptional changes in Latham roots infected with *P. rubi* at 7 dpi. **A.** Variation between infected and control samples of Latham. PCA Plots were generated using average expression of gene level log_2_-CPMs (counts per million). Percentage of variance explained by each principal component (PC) is indicated along the axes. **B.** Volcano plot of differentially expressed genes (DEGs) in control and infected roots of Latham. **C.** Top 16 GO terms associated with the function of DEGs. The GO terms were categorised into Biological Process (BP), Cellular Component (CC) and Molecular Function (MF). A gene was considered differentially expressed when Log_2_FC ≥ 1 with adjusted p-value < 0.05. Gene ratio is the ratio between differentially expressed genes against term size and count represent the number of genes differentially expressed. Data was collected from three biological replicates.

A second biological process significantly overrepresented were 21 DEGs related to the ‘Auxin-activated signalling pathway’ category (**Fig. 2C and Table 1)**. The most highly expressed gene encodes an auxin binding protein (ABP19a) which increased ∼14-fold in *P. rubi* challenged roots when compared to controls. This *RiABP19* gene was previously described in a QTL analysis associated with Latham’s resistance (Graham et al., 2011). The positive association of *RiABP19* to PRR and its upregulation in roots 7 dpi after pathogen challenge prompted further investigation of the function of this gene in host resistance responses.

**Table 1:**
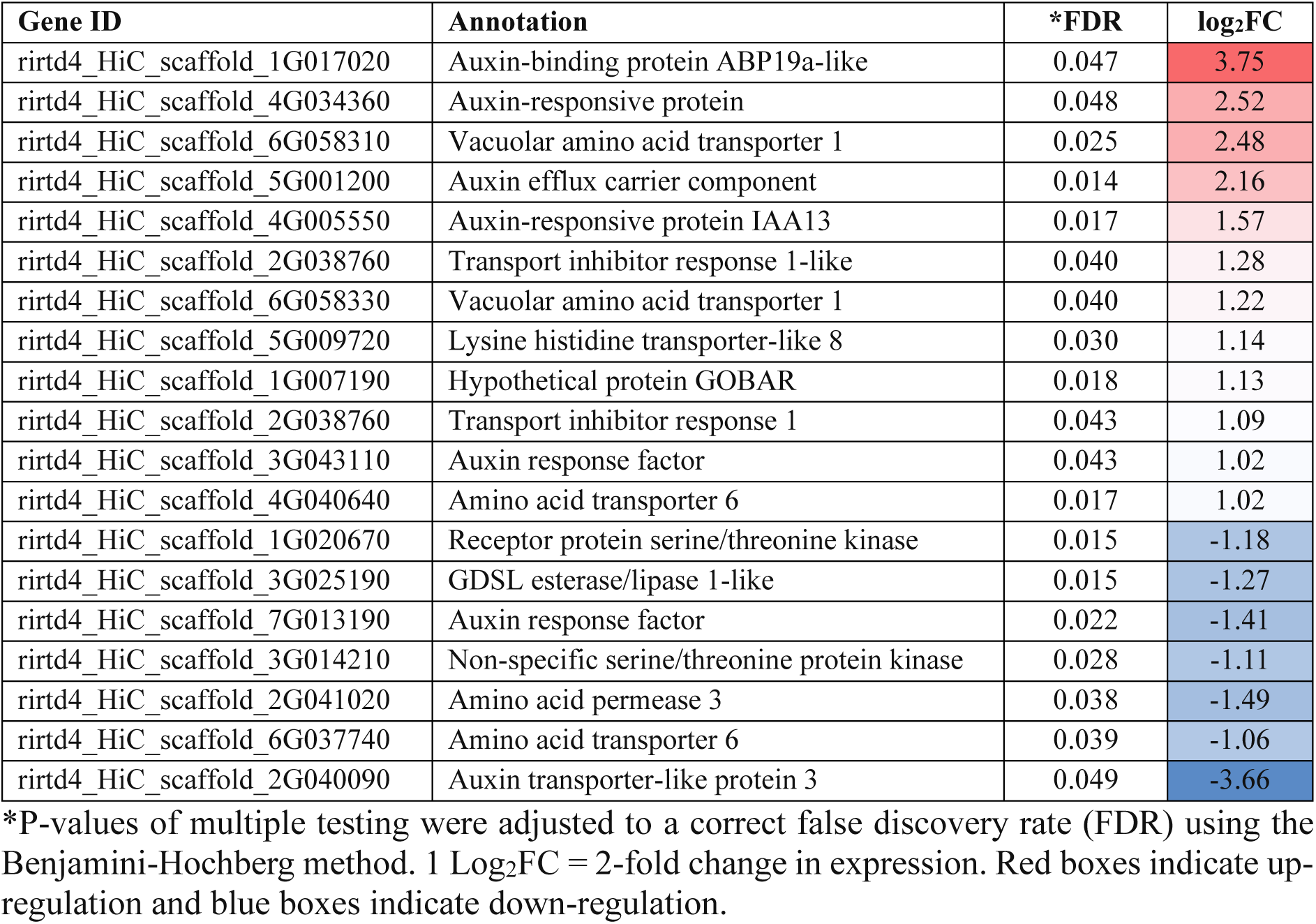
List of differentially expressed auxin-activated signalling pathway genes in Latham roots challenged with *P. rubi*.

### Organisation of multiple copies of *ABP19* genes in the raspberry genome

The cDNA sequence of *RiABP19* obtained from the RNA-seq study of Latham was mapped to chromosome 1 of the red raspberry (cv. Malling Jewel) genome (Price et al., 2023). We identified a cluster of nine *RiABP19* homologues located ∼5Mb upstream of *RiABP19* **(Supplementary Fig. S1)**. These homologues were also mapped to genomic scaffolds of the PRR-resistant cv. Latham. Genes in the cluster were designated as *RiABP19.1-.9* and share 55-95% sequence identity with highly upregulated *RiABP19* **(Supplementary Fig. S2)**. Along with RiABP19 we found *RiABP19.5* and *RiABP19.7* to be expressed in roots. Multiple sequence alignment with previously characterised GLPs/ABP19s highlighted the presence of highly conserved germin BoxA, BoxB, BoxC and KGD motifs (Bernier and Berna, 2001) within RiABP19s **(Fig. 3A)**. A phylogenetic tree comprising sequences from true germins, GLP subfamilies and putative RiABP19s revealed that RiABP19s are members of the GLP subfamily III **(Fig. 3B)**, which is a diverse group including low affinity auxin-binding proteins, expressed in circadian patterns and having roles in disease resistance (Khuri et al., 2001; Mao et al., 2022; Ohmiya, 2002; Pei et al, 2019; Staiger et al., 1999).

**Figure 3:**
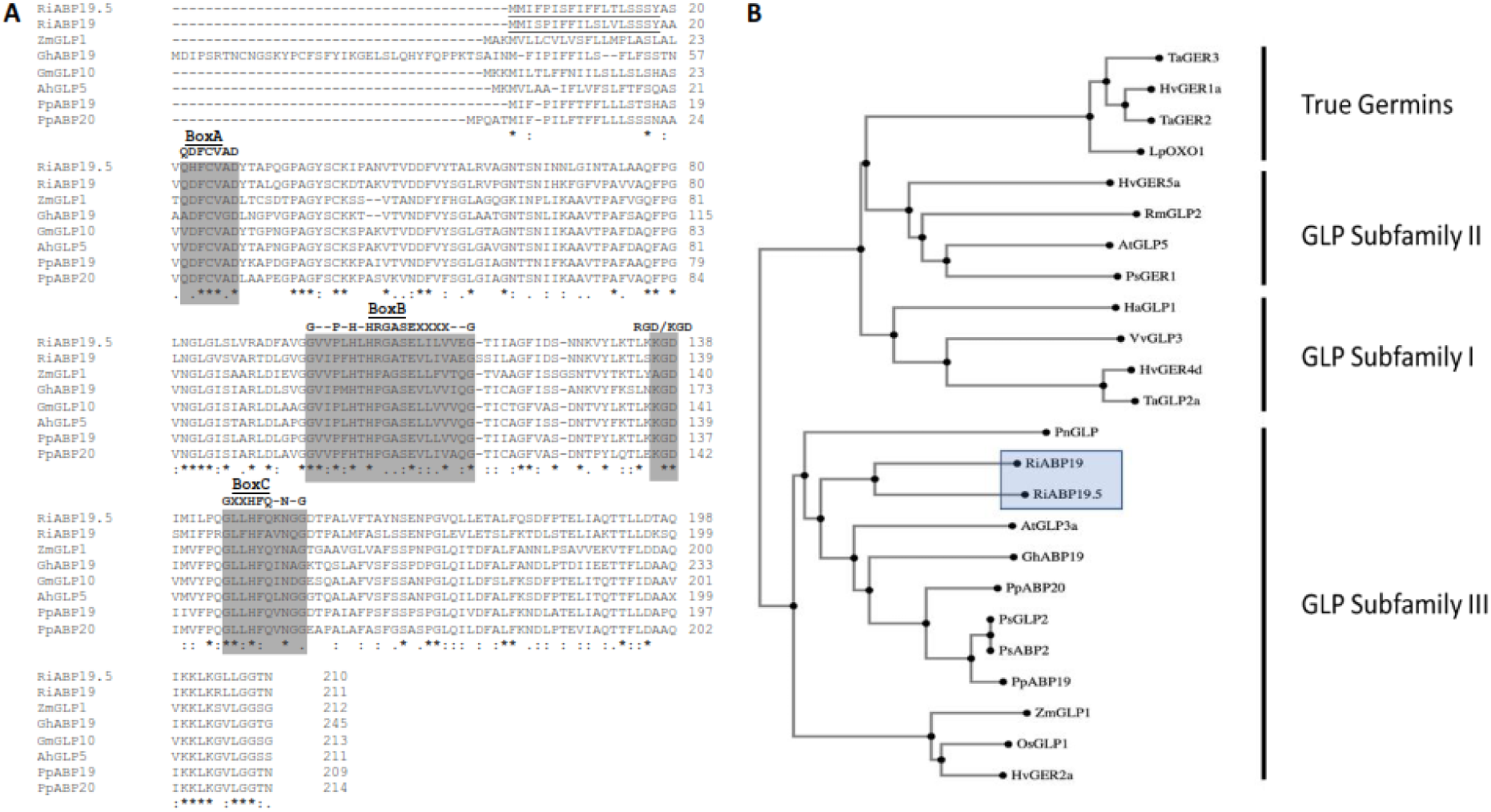
Multiple sequence alignment and phylogenetic analysis of RiABP19 with plant germins and germin-like proteins. **A.** Sequence alignment of raspberry auxin binding proteins with other germin-like proteins. Conserved motifs (Boxes A, B and C) are shaded in grey. X indicates any hydrophobic amino acid. Underlined is the N-terminal signal peptide. Consensus is indicated below the sequences where ‘*’ indicates identical residues, ‘:’ indicates amino acids with similar characteristics and ‘.’ indicates amino acids with similar physiochemical properties. **B.** A phylogenetic tree was created using 24 protein sequences from different plant species. RiABP19s are highlighted in the blue box. Sequences were aligned using Clustal Omega. GeneBank accession numbers of published GLPs used in the analyses are provided in **Supplementary Table S2**.

### RiABP19 is expressed in raspberry roots and upregulated during responses to *P. rubi* challenge in Latham

To examine further expression of *RiABP19, RiABP19.5* and *RiABP19.7*, we used RT-qPCR to compare their expression in leaves and roots in both PRR-resistant and susceptible raspberry cultivars. *RiABP19* expression was significantly higher in roots, whereas *RiABP19.7* showed reduced expression in roots compared to leaves in both cultivars. *RiABP19.5* expression remained consistent in Latham but notably decreased in G. Moy roots (**Fig. 4A**). To evaluate temporal expression of *RiABP19* during pathogen challenge, root samples from Latham were collected 3 h to 23 days after *P. rubi* infection. We found that when compared to uninfected control, *RiABP19* expression increased significantly by 12-fold at 3 h and continued to increase to 26-fold by day 11, subsequently declining to 10-fold by day 14 and thereafter remaining unchanged at day 23 **(Fig. 4B**). Latham showed a clear induction of *RiABP19* expression in response to *P. rubi* challenge at 7 dpi compared to G. Moy where there was no *RiABP19* induction (**Fig. 4C**). Expression of *RiABP19* remained largely unchanged after physical root damage, supporting specific *RiABP19* root induction by *P. rubi*. These expression analyses showed that *RiABP19* expression is specific to raspberry roots and is highly induced in response to infection by *P. rubi,* indicative of a role in the host immune response.

**Figure 4:**
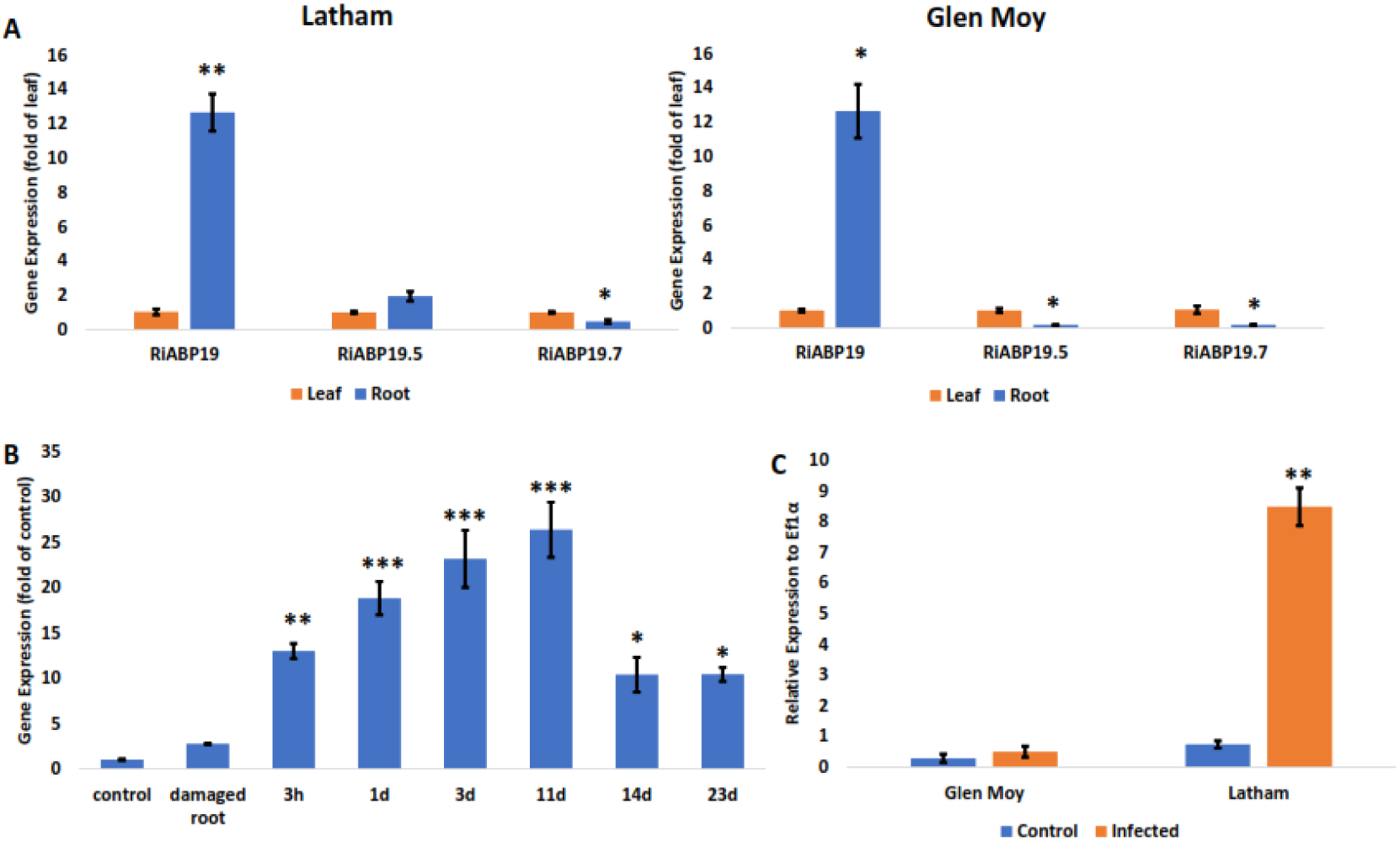
R*i*ABP19 has root-specific expression and demonstrates elevated expression levels when challenged with *P. rubi*. **A.** Expression of three *RiABP19* homologous genes was measured in roots and leaves of PRR-resistant Latham and susceptible G. Moy. Expression in the leaf was set at 1 for each gene. **B.** Expression of *RiABP19* in Latham at 3h, 1, 3, 11, 14 and 23 days after challenge with *P. rubi*. Expression was compared to non-infected control. **C.** Relative expression of *RiABP19* in control and infected roots of Latham and G. Moy, 7 days after *P. rubi* challenge. All expression was relative to the housekeeping gene, Ef1α. Data represents the average of three replicates. Error bars represent standard errors. Significance was calculated by the Student’s t-test and is indicated by asterisks where ***P = <0.001, **P = <0.01 and *P = <0.05.

Sequence scanning of the promoter region 1200 bp upstream of the transcription start site found many auxin-responsive and pathogen response elements (**Table 2 and Supplementary Fig. S3)**. Cis-regulatory elements were identified in both cultivars, but Latham contained two copies of the GT-1 box element and a GCC core element (GCCGCC) associated with defence-related gene expression, 95 bp upstream of the transcription start, which was not present in G. Moy and may account for the lack of defence induced *RiABP19* expression in the susceptible cultivar.

**Table 2:** Auxin-responsive and pathogenesis-related cis-elements in the *RiABP19* promoter.

| Gene | Sequence | Position | G. Moy variation | Description | Reference |
| --- | --- | --- | --- | --- | --- |
| A. Auxin responsive cis-elements |  |  |  |  |  |
| **ASF-1 binding site** | TGACG | 537 (a-3)<br>764 (a-4) | None | Involved in transcriptional activation of several genes by auxin and/or salicylic acid | Liu and Lam, 1994; Niggeweg et al, 2000. |
| **TGA box of AuxRE** | TGACGTAA | 764 (a-4) | None | Found in putative auxin-responsive element (AUXRE) of GH3 promoter | Liu et al., 1994. |
| **ARF binding site** | TGTCTC | 72 (c-1)<br>350 (c-2)<br>564 (c-3) | None | ARF (auxin response factor) binding site found in promoters of early auxin response genes | Ulmasov et al., 1995 |
| B. Pathogenesis-related cis-elements |  |  |  |  |  |
| **W box element** | TTGAC | 110 (a-1)<br>192 (a-2)<br>538 (a-3)<br>765 (a-4)<br>773 (a-5) | None | Recognised by (SA)-induced WRKY DNA binding Proteins in the promoter of NPR1 | Yu et al., 2001. |
| **GT-1 box** | GAAAAA | 549 (b-1)<br>854 (b-2) | One copy in G. Moy | Plays a role in pathogen- and salt-induced gene expression | Park et al., 2004. |
| **SEBF** | YTGTCWC | 72 (c-1)<br>350 (c-2)<br>564 (c-3) | None | Found in the promoter of pathogen-resistant related gene Pr-10a | Boyle and Brisson, 2001. |
| **GCC CORE** | GCCGCC | 95 (d) | Absent in G. Moy | Found in the promoter of many pathogenesis-related genes | Chakravarthy et al., 2003. |

Values in parentheses of the ‘Position’ column represent locations of cis-element annotations in **Supplementary Fig. S3**.

### RiABP19 has structural conservation with true germins

To understand putative function of RiABP19 and whether it may have antifungal properties, deep-learning methodology was applied using IntFOLD-TS utilising integrated LocalColabFold 1.0.0 and trRosetta2 via the IntFOLD7 server to generate tertiary structures predictions and subsequent three-dimensional models and ligand-protein interaction predictions conducted using FunFOLD (Adasme et al., 2021; Anishchenko et al., 2021; McGuffin et al., 2023; Mirdita et al., 2022). Barley germin, PDB:1FI2 was overlayed with the rank 1 RiABP19 model (**Fig. 5A)**. Model quality estimates of folding and interfaces of rank 1 RiABP19 monomer were conducted using ModFOLDdock producing a global model quality score of 0.7194 (scale of 0-1), p = < 0.001 and a confidence score of 4.884E-5. Independent assessment using DeepUMQA produced a global local distance difference test (lDDT) score of 88.67 with only the Nʹ 42 aa residues presenting an lDDT below 80 (**Supplementary Fig. S4)**. This region forms a single α-helix and unstructured region not involved in formation of the ligand-binding region that itself has an lDDT > 80. Assessment with SignalP 6.0 (Teufel et al., 2022) identified this region as containing a 19 aa long signal peptide with a probability of 0.99 and predicted cleavage between 19-20 aa **(Supplementary Fig. S5)**. FunFOLD ligand-binding prediction was assessed using Protein-Ligand Interaction Profiler (PLIP) (Adasme et al., 2021) and the output is shown in **Fig. 5A** inset, where a highly conserved active site consisting of three histidines (His102, His104, His149) and one glutamate (E109) binding to a metal ion within a β-barrel structure was observed.

**Figure 5.**
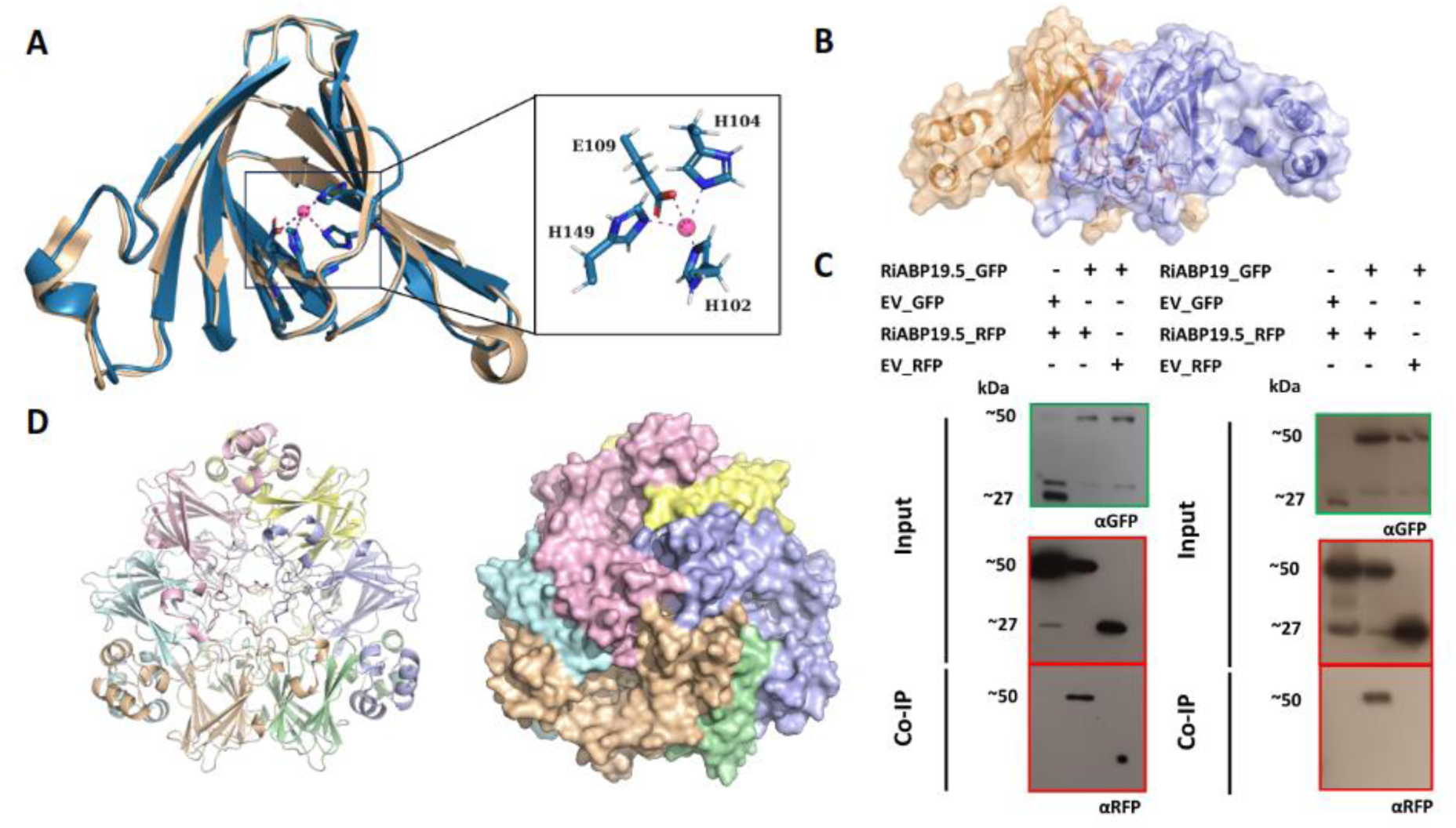
Modelling of RiABP19 and Co-IP assays showing dimerization. **A.** RiABP19 (blue) overlaid with the crystal structure of barley germin (gold). The active site constituting amino acids His102, His104, His149 and E109 are shown in the inset. The pink sphere is the manganese ion of the active site. **B.** Structure of the RiABP19 dimer. **C.** Co-IP of protein extracts from agroinfiltrated leaves confirmed that RiABP19.5:RFP interacts with RiABP19.5:GFP forming a homodimer (left) and with RiABP19:GFP forming a heterodimer (right). Expression of constructs is indicated by +. Empty Vector (EV)_GFP and EV_RFP are used as controls. Green and red boxes indicate a GFP and RFP antibody background respectively. **D.** Modelling of the hexameric structures of RiAPP19. Different colours indicate each monomer. Both ribbon and three-dimensional models are shown.

To examine the potential for RiABP19 to form dimers, a homodimer model was predicted with MultiFOLD utilising LocalColabFold 1.0.0 and 1.3.0, model quality assessed using ModFOLDdockR. The top 5 ranked models were resubmitted to LocalColabFold as template files and reprocessed to produce iteratively refined models using the MultiFOLD server to produce the refined rank 1 model presented in **Fig. 5B**. To test this modelling, Co-immunoprecipitation (Co-IP) assays were conducted. RiABP19 was cloned with a C-terminal GFP fusion (RiABP19:GFP), and RiABP19.5 was cloned with two different C-terminal fusions: one with RFP (RiABP19.5:RFP) and another with GFP (RiABP19.5:GFP). We performed Co-IP using Agrobacterium-mediated transient expression of fused proteins in *N. benthamiana* leaves. Protein extracts from agroinfiltrated leaves and incubation of samples with GFP-trap beads confirmed that RiABP19.5:RFP interacts with RiABP19.5:GFP to form a homodimer and with RiABP19:GFP to form a heterodimer showing that RiABP19 has the potential to form a multimeric protein complex (**Fig. 5C**). We also found that RiABP19 can be predicted to form a hexameric structure similar to the true germin hexamer (PDB:1FI2) (Woo et al., 2000). A hexameric model was predicted by MultiFOLD using the aforementioned methodology producing a rank 1 model with a plDDT of 0.987 and pTM of 0.978 (**Figure 5D**). Model assessment using DeepUMQA returned a global lDDT of 81.38 (**Supplementary Fig. S6)** with multiple residues involved in protein-protein interactions forming the complex (**Supplementary Table S3**). These predictions show that RiABP19 has structural conservation with barley germin and may have similar functional roles.

### *NbABP19* is a positive regulator of plant immune responses in *Nicotiana benthamiana*

As raspberry genetic modification is not accessible, we used VIGS to silence *RiABP19* orthologs in the model plant, *N. benthamiana*; a host for a related Phytophthora species. *RiABP19* was used to search the Sol Genomic database (https://solgenomics.net/) for orthologs in *N. benthamiana*, identifying four orthologs with 52-61% identity. Based on sequence identity these four ortholog existed as pairs, *NbABP19a/b* and *NbABP19c/d* (**Supplementary Fig. S7A**). Avoiding the conserved cupin domain, divergent sequences at the terminal 5’ or 3’ regions from each gene pair were selected for antisense cloning into Agrobacterium-delivered tobacco rattle virus vector (TRV) for transient VIGS in the plant (Gilroy et al., 2007). Three vectors were constructed to silence *NbABP19a/b*, *NbABP19c/d* and all four *NbABP19* genes (**Supplementary Fig. S7B**). All experiments used control plants generated by inoculating Agrobacterium with the vector used to transiently express the sense copy of full length GFP. Significant silencing of *NbABP19a/b*, *NbABP19c/d* and all four *NbABP19* genes was confirmed by RT-qPCR after three weeks of VIGS infiltration (**Supplementary Fig. S7C**).

To investigate the effect of *NbABP19* silencing on host immune responses, cell death (CD) assays were conducted 2-3 weeks post inoculation of the TRV constructs either expressing antisense fragments of *Nbabp19* or control expressing GFP. Leaves of VIGS plants were inoculated with Agrobacterium transiently expressing pathogen proteins and combinations of INF1, Cf4/Avr4 and Pto^Y207D^ to induce immune regulated CDs and were counted at 7 dpi. INF1 is a *P. infestans* elicitor protein that triggers Pathogen Triggered Immunity (PTI) in *N. benthamiana* plants (Kamoun et al., 1998). Avr4 is an apoplastic virulence protein secreted by tomato fungal pathogen *Cladosporium fulvum* which triggers a hypersensitive response (HR) when co-infiltrated with its cognate tomato host receptor Cf4 (Van der Hoorn et al., 2000). Although there was no difference in INF1 CDs, the HR to Cf4/Avr4 was significantly lower in *Nbabp19* silenced plants, particularly in plants where all four genes were silenced **(Fig. 6A)**. Pto^Y207D^ is an autoactive version of protein kinase Pto which normally functions in recognition of the *Pseudomonas syringae* virulence protein AvrPto (King et al., 2014). Results of assays conducted to examine whether *NbABP19* silencing affects Pto^Y207D^-induced HR, showed a significant decrease in HR **(Fig. 6A)**, suggesting involvement of NbABP19 in SGT1 dependent, effector-triggered immunity (ETI) (Gabriels et al., 2007).

**Figure 6:**
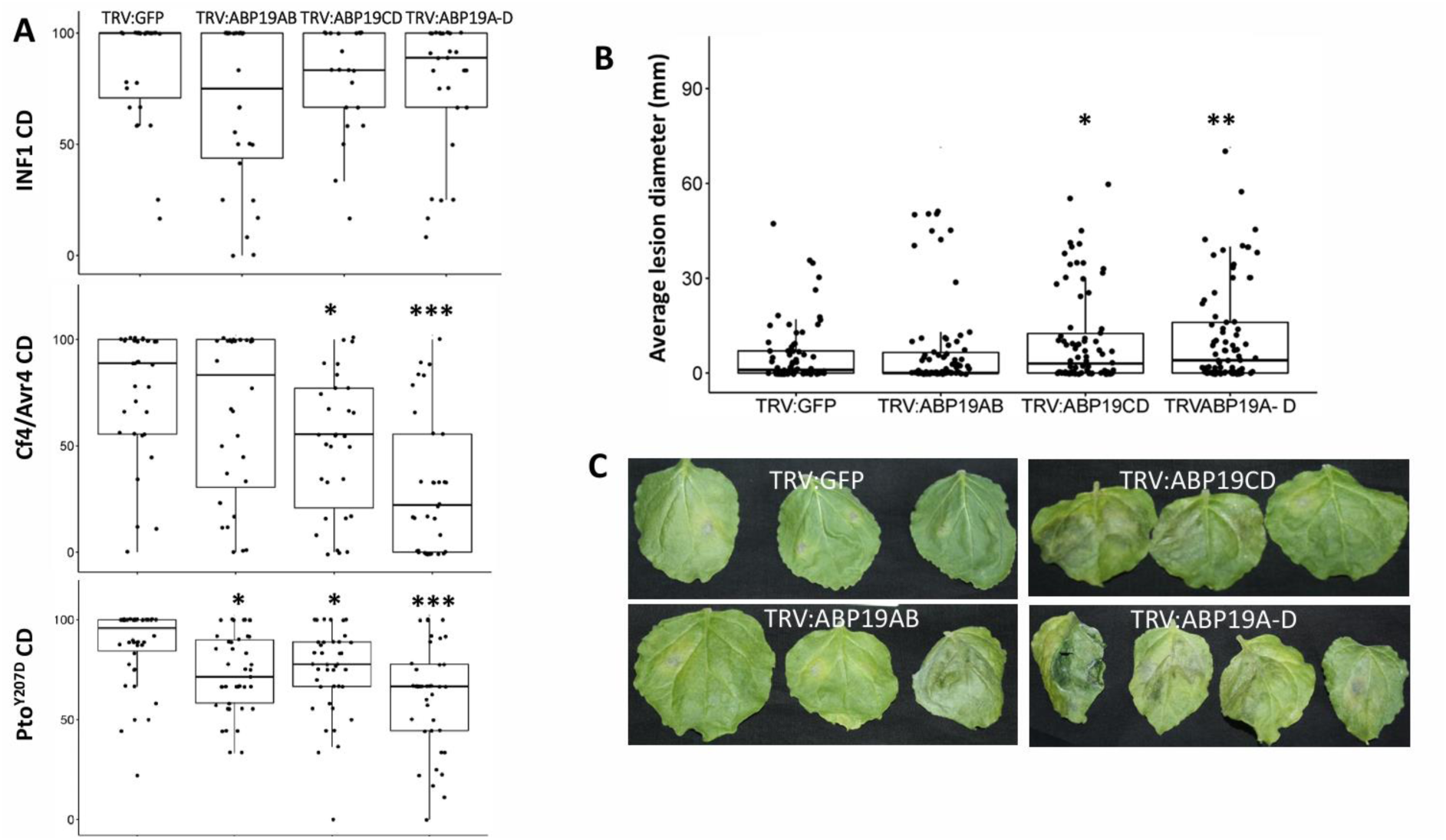
Silencing of NbABP19 leads to a reduced defence response and increased susceptibility. **A.** Effect of NbABP19 silencing on different immune cell deaths (CDs). Box plots depict average percentage of infiltrated sites that developed CD at 5 dpi for autoactive PtoY207D and 7dpi for Cf4/Avr4 and INF1. Plots represent combined data from five biological replicates. **B.** Effect of NbABP19 silencing on *P. infestans* colonization. Detached leaves of NbABP19 silenced plants were inoculated with *P. infestans* spores. Box plots depict average lesion diameter. Data represents the combination of four independent experiments (number of leaves, n = >50). **C.** Photos of *P. infestans* infection on leaves expressing different constructs. Error bars are standard errors and significant differences to the GFP control were calculated by the Holm-Sidak method, one-way ANOVA indicated by asterisks, * P = <0.05, **P = <0.01 and *** P = <0.001.

To investigate the impact of *NbABP19* silencing on *P. infestans* colonization, detached leaves from *N. benthamiana* plants three weeks after inoculation with the different VIGS constructs were infected with *P. infestans* zoospore suspension. After 12 days post-inoculation, the number of infected leaves were compared to plants expressing the *TRV:GFP* control. Results showed that leaves silenced for all four copies of *NbABP19* using *TRV:ABP19A-D* were more susceptible to *P. infestans* infection **(Fig. 6B and C)**. Moreover, both necrosis and sporulation of *P. infestans* were higher in *NbABP19* silenced plants, with percentage being highest in *TRV:ABP19A-D* plants **(Supplementary Fig. S8)**. Taken together our results suggest that *NbABP19* functions as a positive regulator of plant immunity as silencing all four copies in *N. benthamiana* plants perturbs the HR induced by Avr4/Cf4 and increases susceptibility to *P. infestans* infection.

## Discussion

Root anatomy and architecture play an important role in how plants respond to various biotic and abiotic stresses. Vigorous root systems like cv. Latham can provide a greater interface for interaction with a wide range of microbes in soil and allow greater opportunities for pathogens to attach to and attack their host. On the other hand, plants with weaker root systems like cv. G. Moy may provide fewer niches to colonise but may be more amenable to microbial interactions in a trade-off for better nutrient acquisition and water capture. Radial movement of pathogens through root cells towards the vascular bundle is a key factor in successfully colonising roots. RNA-seq analysis of Latham challenged with *P. rubi* revealed upregulation of multiple pathogen response genes suggesting an active defence response has been triggered (**Fig. 2**). However, cross-sectional analysis of the primary root of our selected raspberry cultivars showed that Latham possesses a thicker cortex than G. Moy, which could be responsible for delaying pathogen progression (**Fig. 1C**). Studies in tomatoes indicated progression of *Ralstonia solanacearum* was delayed in the cortex of resistant roots but advanced more rapidly in susceptible roots (Caldwell et al., 2017). Thicker cortex and expression of disease response genes in Latham indicate a cultivar able to inhibit pathogen progression and continue to produce, in line with previous studies, new healthy roots following *P. rubi* infection (**Fig. 1B**). This contrasts with the simpler root architecture of G. Moy and quicker disease progression, with roots darkening with root rot symptoms seven days post-infection (**Fig. 1A**). Collectively, these findings suggest that robust root growth and structural difference in root cortex may be components of host tolerance to PRR in Latham.

Transcriptional changes taking place in Latham challenged by *P. rubi* identified the auxin-activated signalling pathway among the top 16 GO categories (**Fig. 2**). Auxin has emerged as a key player in plant immunity, orchestrating responses to pathogenic challenges (Gilroy and Breen, 2022). Considering the importance of auxin in both root development and disease progression, combined with our focus on resistance to PRR, we prioritised characterisation of a GLP gene, RiABP19, which was significantly up-regulated specifically in PRR-resistant raspberry cultivar Latham in response to *P. rubi* infection (**Fig. 4C and Table 1).** There is increasing evidence for involvement of GLPs in host resistance (Govindan et al., 2024). For example, cotton GhABP19 not only displays regulation of host resistance, but recombinant purified protein also displayed antifungal activity (Pei et al., 2019). Studies have shown that GLP gene members are abundant in higher plants with 30 GLPs in Arabidopsis, 15 in tomato and 21 in soybean. Some GLP clusters have been described to function cooperatively in plant immune responses; for example, a cluster of 12 GLPs was found to be located within a QTL associated with broad-spectrum disease resistance on chromosome 8 in rice (Manosalva et al., 2009). Genomic analyses of Latham revealed a cluster of nine *RiABP19* homologs (*RiABP19.1-RiABP19.9*) located ∼5Mb upstream of *RiABP19* (**Supplementary Fig. S1**). Nine of 10 raspberry RiABP19s shared significant homology and contained conserved GLP protein motifs (**Supplementary Fig. S2**). Sequence alignment with other published GLPs and phylogenetic tree analysis grouped RiABP19 members within the GLP subfamily III (**Fig. S3**), which is a heterogeneous group with members exhibiting different functions including proteins involved in auxin metabolism and disease resistance (Khuri et al., 2001; Pei et al., 2019).

Further gene expression analysis of *RiABP19* and detectable homologs *RiABP19.5* and *RiABP19.7* revealed expression in raspberry leaves and roots, with expression of *RiABP19* being 10-fold greater in roots than in leaves (**Fig. 4A**). GLPs are expressed in different tissues, at different developmental stages and under different conditions (Barman and Banerjee, 2015). Examination of temporal expression of *RiABP19* after challenge of Latham with *P. rubi* zoospores by RT-qPCR revealed a 12-fold induction by 3 hpi which steadily increased until 11 dpi when compared to uninfected control (**Fig. 4B)**. As *RiABP19* is induced in Latham after pathogen challenge from as early as 3 hpi, we can speculate that RiABP19 plays a role in inducible immune responses in Latham roots. Analysis of the promoter regions of *RiABP19* revealed the presence of multiple auxin and pathogenesis-related cis-elements (**Table 2**). The D4 element of the GH3 gene, which is important for auxin inducibility (Ulmasov et al., 1995) was present in three positions. This element was also present in the promoter region of peach *PpABP19*, which was shown to bind auxin (Ohmiya, 2002). Among four pathogen resistance-related elements found in the promoter of *RiABP19*, three (W-box, GT-1 box and SEBF) were also present in the *GhABP19* promoter, the protein of which is involved in fungal resistance in cotton (Pei et al., 2019). A GCC core element (GCCGCC) associated with defence-related gene expression and present in many pathogen-responsive genes (Brown et al., 2003) was present in the promoter of Latham but was absent in G. Moy which may support differential regulation of *RiABP19* in different cultivars (**Fig. 4C**). Overall, our findings highlight that auxin-responsive cis elements exhibited similarity in their presence and location in promoters of Latham and G. Moy RiABP19 whereas, pathogen resistance-related cis-elements showed presence/absence and abundance differences.

Modelling of plant protein structure can be successfully applied to understand the function of a protein (Rasheed et al., 2020). Three-dimensional model of RiABP19 showed homology to barley germin, where the substrate binding site consisting of a manganese ion bound by three histidine and one glutamine residue are highly conserved (**Fig. 5A**). Site-directed mutagenesis of this active site in rice *OsRGLP1* decreased or lacked SOD activity and was associated with auxin-binding activity in maize ZmABP1 (Shahwar et al., 2023). GERs and GLPs are known to form homodimers, trimers, and hexamers (Dunwell et al., 2001). In Arabidopsis, it has been reported that two GLPs, PDGLP1 and PDGLP2, form a complex to control primary root growth (Ham et al., 2012). Pull-down and Co-IP assays here showed that RiABP19.5 can interact to form complexes of homo- and heterodimeric proteins which may be important for its function. Using protein modelling, we also predict that RiABP19 can assemble into a hexameric structure similar to the barley germin homohexamer, which is important for SOD and OXO activity (**Fig. 5D**) and would require confirmation using Cryo-EM techniques (Woo et al., 2000).

Due to the lack of agrobacterium-mediated transformation, gene editing or VIGS systems in raspberry, functional characterisation of ABP19 in resistance was transferred to a well-characterised alternative pathosystem, host *N. benthamiana* and pathogen *P. infestans* (McLellan et al., 2013). Four endogenous *NbABP19* orthologs of *RiABP19* were identified in *N. benthamiana* and these were silenced in pairs (*TRV:ABP19AB* and *TRV:ABP19CD*), and all four *NbABP19* genes altogether (*TRV:ABP19A-D*). Results showed that silencing either pair of homologs alone did not have a significant effect on disease development but when all four genes were silenced together we observed significantly increased susceptibility to *P. infestans* and increased pathogen sporulation when compared to plants inoculated with *TRV:GFP* control (**Fig. 6B and Fig. 6C and Supplementary Fig. S8**). Silencing of all four *NbABP19* genes also perturbed CD responses to previously characterised ETI triggers in *N. benthamiana* (Cf4/Avr4 and PTO^Y207D^) (**Fig. 6A**). INF1-CD was also examined but no significant difference was observed (data not shown). Our data supports that ABP19 is a positive regulator of defence, involved in signalling downstream of more than one immune pathway. Notably, both Cf4/Avr4 & Pto^Y207D^ HRs share components of defence signalling including SOBIR1, NRC1 and a MAPKKKα gene (del Pozzo et al 2004). The RiABP19 results indicate a function as a positive regulator in immune signalling pathways targeted by Phytophthora effector proteins.

In conclusion, our study indicates an important role for RiABP19 in host resistance in raspberries. Further functional analysis by overexpression and gene editing will determine *RiABP19* as a suitable candidate for developing PRR-resistant raspberry cultivars.

## Materials and Methods

### Plant growth and sampling

Raspberry plants were grown in a glasshouse with a temperature of 20°C and a lighting cycle of 16 h followed by 8 h at 18°C without artificial lighting. For RNA-sequencing and RT-qPCR, samples were collected from eight-week-old healthy Glen Moy and Latham plants removed from their pots and transferred to Erlenmeyer flasks containing water, with or without *Phytophthora rubi* inoculum. Seven days after pathogen challenge, around 1-2cm of the root tips of three independent biological samples were collected and stored at -80°C. For the remaining experiments, samples were collected from hydroponically propagated G. Moy and Latham plants. Softwood with a clear stem and 2-3 internodes were dipped in rooting hormone gel, Clonex (Growth Technology) and placed in autoclaved sand equipped with a mist unit. After rooting they were transferred to Nutrient Film Technique hydroponic tanks filled with weak nutrient solution (1g/L of Solufeed pH 6-6.5).

*Nicotiana benthamiana* plants were cultivated in pots with general-purpose compost in a glasshouse at 22°C for 16 h, followed by 8 h at 18°C, with 130-150 μE m^-2^ s^-1^ light intensity and 40% humidity. Plants aged four to five weeks were utilised for transient expression, cell death assays, and *Phytophthora infestans* infection assays.

### Raspberry root infection with *Phytophthora rubi*

*P. rubi* isolate, SCRP333, was cultured on rye agar media supplemented with 100μg/mL ampicillin. After two weeks of growth, small plugs from actively growing regions were covered with soil solution prepared by soaking 150g general glasshouse compost in 1.5L distilled water (DW). The solution was stirred for 1h at room temperature and then passed through a double layer of filter paper twice. The plugs with soil solution were covered with aluminium foil, incubated at 15°C and changed three times over 3-4 days. After confirmation of sporangia development, plugs were placed in a 4°C fridge for 30 min to shock the sporangia and release zoospores. Additionally, 50mL sterile Petri’s solution (0.074g KCL, 0.472g Ca(NO_3_)_2_, 0.296g MgSO_4_ and 0.136g KH_2_PO_4_ diluted in 1L DW and autoclaved) was added to encourage zoospore release. This inoculum was used to infect raspberry roots in soil-free media.

Raspberry cuttings with an established root system were washed with DW and transferred to Erlenmeyer flasks with 200mL DW and 5mL inoculum. Flasks were covered and kept at 18^ο^C, to facilitate infection. Leaves were sprayed daily with DW. Root imaging and samples were collected after seven days.

### RNA extraction and sequencing

Total RNA extraction of root tips was carried out using a Macherey-Nagel NucleoSpin RNA Plant mini kit. It included an equal volume of Plant Isolation Aid (Invitrogen) and centrifugation before RNA isolation. Total RNA was quality checked using a Bioanalyzer 2100 (Agilent) to ensure a RIN >7. In total, 1μg RNA per sample was used as input material for sample preparations. Strand-specific dUTP libraries and Illumina paired-end 150 bp sequencing were completed by Novogene (UK) Company Limited.

### Assembly of raspberry RiRTD4 from Illumina short-read RNA-seq

Raw RNA-seq short-read derived from four publicly available datasets (PRJNA476755, PRJEB28528, PRJNA354231, PRJNA560307) and four in-house datasets including this dataset (PRJEB73351) was used to assemble a reference transcript dataset (RTD) for raspberry (**Supplementary Table S4**). RNA-seq reads were processed using Fastp v0.20.1 to remove adapters and filter reads with a quality score of < 20 and a length of < 30 bases (Chen et al., 2004). STAR v2.7. was applied to map trimmed reads to the reference cv. Anitra genome with an allowance for a 2% mismatch across reads for variation between cultivars (Davik et al., 2022; Dobin and Gingeras, 2015). Stringtie v2.1.5 (Pertea et al., 2015) and Scallop v0.10.5 Shao and Kingsford, 2017) was used to construct both unstranded and stranded transcript assemblies. RTDmaker (https://github.com/anonconda/RTDmaker) was then used to merge assemblies across samples and filter out low-quality transcripts. Transcripts showing non-canonical splice junctions were removed as were transcripts with splice junctions supported by fewer than 10 reads in fewer than two samples. Fragmentary transcripts whose length was < 70% of the gene length, transcripts with low abundance with expression levels < 1 transcript per million reads (TPM) in fewer than two samples, and mono exonic-antisense fragment transcripts located on the opposite strand of the gene with a length < 50% of the predicted gene length were all removed. Without access to long-read sequence data, we padded shorter 5′ and 3′ transcript ends of individual genes to 5′ and 3′ ends of the longest gene transcript to improve quantifications as described previously (Rapazote-Flores et al., 2019) (**Supplementary Fig. S9**).

### Draft genome assembly for Latham

A draft genome was previously established for G. Moy (Hackett et al., 2018). We used the same protocols to extract genomic DNA from young leaves to fragment and construct libraries. Sequencing was performed as two independent runs on a MiSeq (Illumina) using paired-end 2 × 300 bp v3 kits (Illumina) and CLC_assembler from the CLCBio suite (v 4.10.86742, with a range of k-mer sizes). Sequencing was done in the absence of mate pairing. This gave a draft genome for Latham consisting of 193,970 output scaffolds covering 271 Mbp of an estimated 300 Mbp genome.

### Differential gene expression analysis

RNA-seq reads from Latham and G. Moy roots challenged with *P. rubi* were mapped onto RiRTD4 using Salmon (Patro et al., 2017). Differential gene expression analysis was performed using the 3D RNA-seq pipeline (Guo et al., 2021). Transcript-level abundances (TPM) from each Salmon file, estimated counts, and effective lengths were imported into 3D RNA-seq. Samples were filtered to remove poorly expressed transcripts (≥ 1 CPM in ≥ 1 samples and data normalised gene and transcript read counts to log_2_-CPM using weighted trimmed mean of M-values (Bullard et al., 2010). Limma R package was used for 3D RNA-seq expression comparisons. To compare expression changes between experimental design conditions, contrast groups were set as Control versus Infected. For DE genes/transcripts, the log_2_ fold change of gene abundance was calculated based on contrast groups and the significance of expression changes was determined using a t-test. P-values of multiple testing were adjusted with Benjamini-Hochberg to correct the false discovery rate (FDR). A gene/transcript was significantly DE in a contrast group if it had adjusted p-value < 0.05 and Log_2_FC ≥ 1.

Gene ontology (GO) annotations were identified by sequence similarity to strawberry and GO enrichment for differentially expressed genes was performed using g:Profiler (Raudvere et al., 2019).

### Quantitative RT-PCR gene expression analysis

cDNA was reverse transcribed from 800ng-1μg of total RNA using Takara RNA to cDNA EcoDry Premix (Takara Bio) as per manufacturer instructions. RT-qPCR was performed using PowerUp™ SYBR™ Green Master Mix (Thermo Fisher Scientific). Reactions were carried out in a StepOnePlus™ Real-Time PCR System machine. Reactions were incubated at 50°C for two min followed by an initial denaturation at 95°C for 15 min, before 40 cycles of 95°C for 15 sec, and 60°C for one min. The Pffafl method (Pfaffl, 2001) as used to calculate fold changes relative to control plants. Elongation factor 1α (Ef1α) of *R. idaeus* was used as a housekeeping gene. Primer sequences are listed in **Supplementary Table S5**.

### Sequence analysis

The open reading frame (ORF) of *RiABP19* and homologues were identified using ExPasy (Gasteiger et al., 2003). Multiple sequence alignment and phylogenetic tree analysis were carried out using Clustal Omega with default parameters (Madeira et al., 2022). To identify cis-regulatory elements in the promoter region of *RiABP19*, 1200bp upstream genomic sequence from the transcriptional start site of Latham and G. Moy *RiABP19* was selected and scanned using NEW PLACE (www.dna.affrc.go.jp/PLACE) (Higo et al., 1999).

### Protein modelling

Tertiary and quaternary structures, as well as protein-ligand interactions of RiABP19.10, were predicted and refined using the latest versions of IntFOLD-TS, MultiFOLD, FunFOLD as part of the IntFOLD7 suite of modelling tools (McGuffin et al., 2023; Adiyaman et al., 2023; Roche et al., 2013), taking advantage of recently integrated 3D modelling with both trRosetta2 (Anishchenko et al., 2021) and LocalColabFold (Mirdita et al., 2022). Structures were subjected to further refinement using ReFOLD (Adiyaman et al., 2023) and model quality assessment using ModFOLDdock, ModFOLDdockR (Edmunds et al., 2023) and DeepUMQA (Guo et al., 2022). FunFOLD ligand-binding prediction was assessed using the Protein-Ligand Interaction Profiler (PLIP) (Adasme et al., 2021).

### Generation of protein fusion clones and VIGS constructs

Full-length *RiABP19* and *RiABP19.5* were amplified from Latham cDNA using gene-specific primers containing attB recombination sites for the Gateway cloning system (Invitrogen, UK). The stop codon was removed to prevent translation termination before the C-terminal reporter. Resulting PCR products were purified and cloned into pDONR201 utilising BP clonase^®^ II and subsequently inserted into pB7FWG2 for GFP fusion and pK7RWG2 for RFP fusion using LR clonase^®^ II. Cloned proteins were transferred into the *Agrobacterium tumefaciens* strain, GV3101 and used for transient assays.

*NbABP19a/b* and *NbABP19c/d* silencing trigger regions were amplified from *N. benthamiana* DNA via standard amplification PCR, using KOD Hot Start DNA Polymerase (Novagen). The hybrid *NbABP19a-d* fragment was obtained using overlap-extension PCR to combine *NbABP19a/b* and *NbABP19c/d* PCR products. Three VIGS PCR products were cloned into pBinary Tobacco Rattle Virus (TRV) vectors (Liu et al, 2002), between HpaI and EcoRI sites in antisense orientation. As a control, TRV expressing GFP was used (Gilroy et al., 2007). Four-leaf-stage wild-type *N. benthamiana* were agroinfiltrated at an OD_600_ of 0.25 for each VIGS construct and 0.1 for RNA1 (McLellan et al., 2013). Primer sequences are provided in **Supplementary Table S5.**

### Agrobacterium-mediated transient expression

Agrobacterium strains with cloned products were grown in YEB media supplemented with appropriate antibiotics and incubated at 28^ο^C overnight. Cultures were spun at 4,500g and the pellet was resuspended in sterile 10 mM MES and 10 mM MgCl_2_ buffer supplemented with 200 μM acetosyringone. Final optical density (OD_600_) was measured and adjusted as desired. Four to five weeks old *N. benthamiana* plants were used for infiltration. The abaxial side of leaves was pressure infiltrated using a 1mL syringe after wounding with a needle.

### Infection assays

*N. benthamiana* plants were infiltrated with agroinfiltration buffer containing pathogenic proteins INF1, Cf4/Avr4 and PTO^Y207D^ at an OD_600_ of 0.5 for each construct except INF1 (OD_600_ = 0.4). Three to four leaves on six plants were infiltrated for each construct. CDs were counted after 7 dpi and recorded as described in (Gilroy et al., 2011). *P. infestans* infection assays on *N. benthamiana* VIGS plants were conducted with *P. infestans* strain 88069 (McLellan et al., 2013). Statistical analysis was conducted using one-way ANOVA (Holm-Sidak method) or Student’s t-test.

### Co-immunoprecipitation

The three largest middle leaves of *N. benthamiana* plants were infiltrated with agroinfiltration buffer containing the fusion protein constructs at an OD_600_ of 0.5. Leaf tissues were collected after 48 h, and proteins were extracted (McLellan et al., 2013). Immunoprecipitation of GFP-tagged protein fusions was performed using GFP-Trap® Magnetic Agarose beads (ChromoTek). Resulting samples were separated by PAGE and Western blotted (McLellan et al., 2013). Immunoprecipitated GFP fusions and coimmunoprecipitated RFP fusions were detected using appropriate antibodies.

## Data Availability

The RNA-seq data was deposited in the European Nucleotide Archive under accession number PRJEB73351. The Latham sequence was assembled into scaffolds and is available from the authors. The assembled transcriptome RiRTD4 can be made available on request. AlphaFold models can also be made available on request. All other relevant data are presented within the paper and its supplementary files.

## Author Contribution

MG and EG, conceived the study. Plants and *P. rubi* were grown and measurements taken by RO, KS and EG. Gene silencing and co-immunoprecipitation experiments were performed by RO, CEO, LW, AB, HM and EG. Gene expression analysis was performed by RO, CS and JF. Genome and transcriptome sequence assemblies were prepared by LM. Protein folding analysis was performed by RF. The paper was written by RO, EG, CS, RF and MG. All the authors have reviewed and approved the manuscript.

## Supplementary information

**Supplementary Figure 1:** Distribution of the RiABP19 homologues on chromosome 1 of red raspberry (cv. Malling Jewel).

**Supplementary Figure 2:** Protein sequence alignment of RiABP19 and homologues located on chromosome 1 of red raspberry.

**Supplementary Figure 3:** Sequence of 1200 bp promoter region of *RiABP19* in Latham and G. Moy.

**Supplementary Figure 4:** Result of model quality assessment of RiABP19 monomer using the DeepUMQA method.

**Supplementary Figure 5:** SignalP-6.0 prediction of RiABP19.

**Supplementary Figure 6:** Model quality assessment of RiABP19 hexamer using the DeepUMQA model.

**Supplementary Figure 7:** Virus-induced gene silencing of *NbABP19s* in *Nicotiana benthamiana* plants.

**Supplementary Figure 8:** Silencing of *NbABP19* results in increased susceptibility to *Phytophthora infestans* infection.

**Supplementary Figure 9:** Workflow of the pipeline used to develop reference transcript dataset (RTD).

**Supplementary Table 1:** List of differentially expressed genes (DEGs) categorised under ‘defence response’ in challenged Latham roots at 7 dpi.

**Supplementary Table 2:** Characterised GLPs with accession numbers that were used to run sequence alignment and phylogenetic tree analysis.

**Supplementary Table 3:** Interface residue lDDT of homohexamer model of RiABP19.

**Supplementary Table 4:** Public and in-house RNA-seq datasets used to assemble a raspberry reference transcript database.

**Supplementary Table 5:** Primer sequences used for cloning and RT-qPCR.

