## Supplemental Figures and Tables for "Transcriptional response to Phytophthora root rot in raspberry identifies *RiABP19*, a Germin-like protein (GLP) gene with a putative role in resistance"

**Raisa Osama et al**

**Supplementary Documents**

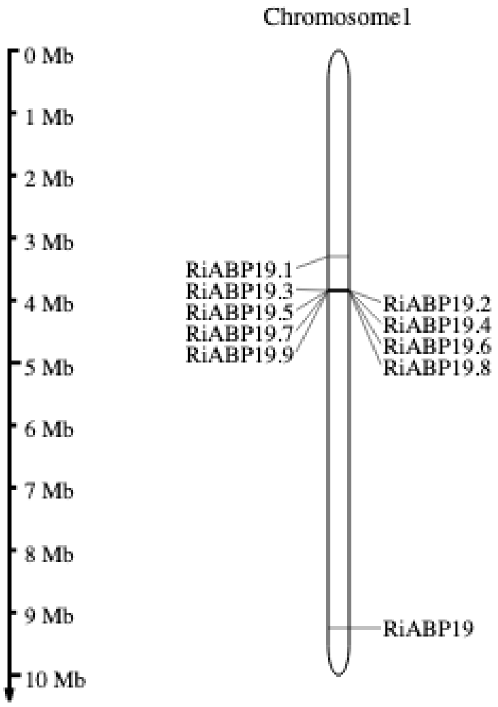


**Supplementary Figure S1:** Distribution of the *RiABP19* homologues on chromosome 1 of red raspberry (cv. Malling Jewel). A cluster of nine *RiABP19* homologues (designated as *RiABP19.1*-*RiABP19.9*) was found to be located ~5Mb upstream of the highly upregulated *RiABP19* identified in Latham roots challenged with *P. rubi* at 7 dpi.

```
RiABP19         MMISPIFFILSLVLSSSYAAVQDFCVADYTALQGPAGYSCKDTAKVTVDDFVYSGLRVPG 60
RiABP19.8       MMIFPIFFTCFLLLSSSHASVQDFCVADFTAPEGPAGYSCKKPAKVTVDDFVFSGLGIAG 60
RiABP19.9       -MIFPIFFAFVLLLSTSHAAVQDFCVADYQAPEGPAGYSCKKPAKVTVDDFVFSGLGVAG 59
RiABP19.1       MITFPVLFVFCLIFSSSYAAVQDFCVADYTAPQSPAGYFCKNPANVTVDDFVYTGLEVAG 60
RiABP19.2       MMIFPISFIFFLTLSSSYASVQDFCVADYTAPQGPAGYSCKIPANVTVDDFVYTGLGVAA 60
RiABP19.4       MMIFPISFIFFLTLSSSYASVQDFCVADYTAPQGPAGYSCKIPANVMVDDFVYTALRVAG 60
RiABP19.5       MMIFPISFTFFLTLASSYASVQDFCVADYTAPQGPAGYSCKIPANVTVDDFVYTALRVAG 60
RiABP19.6       -MMFPISFIFFLTLASSYASVQDFCVADYTAPQGPAGYSCKIPANVTVDDFVYTALRVAG 59
RiABP19.3       ------------------------------------------------------------ 0
RiABP19.7       MMKLPILFVFSLILSSSYAAVQDFCVADYTAPQGPAGYSCKNPENVTVDDFVYSALGVPG 60

RiABP19         NTSNIHKFGFVPAVVAQFPGLNGLGVSVARTDLGVGGVIPFHTHRGATEVLIVAEGSSIL 120
RiABP19.8       NTTNIIKAAVTPAFAAQFPGVNGLGISLARLDLAVDGVIPFHTHPGASEVLVVVEGT-IC 119
RiABP19.9       NTTNIIKAAVTPAFAAQFPGVNGLGLSLARLDLAAGGVVPFHTHPGASEVLVLVQGT-LV 118
RiABP19.1       NASKINKVRLNPAFVSQFPGLNGLGLSLARADLAVGGAIPLHTHHEASEIILVAEGT-VV 119
RiABP19.2       NTSNINNVGITPAFSSQFPGLNGLGLSLVRADFAVGGVSPIHVHRGASELILVVEGT-II 119
RiABP19.4       NTSNINNLGVTTAFSAQFPGLNGLGLSLVRADFAVGGVSPIHVHRGASELILVVEGT-II 119
RiABP19.5       NTSNINNLGINTALAAQFPGLNGLGLSLVRADFAVGGVVPLHLHRGASELILVVEGT-II 119
RiABP19.6       NTSNINNLGINTALAAQFPGLNGLGLSLVRADFAVGGVSPLHVHRGASELILVVEGT-II 118
RiABP19.3       ----------------------------------------MHTHHGASEIVLVVEGT-II 19
RiABP19.7       NTSNMIKTGITTAFSSQFPGLNGLGLSLARADFAVGGVIPMHTHHGASEIVLVVEGT-II 119
                                                        :* *  *:*::::.:*: :

RiABP19         AGFIDSNNKVYLKTLSKGDSMIFPRGLFHFAVNQGDTPALMFASLSSENPGLEVLETSLF 180
RiABP19.8       AGFVASDNTVYLQTLEQGDSMVFPQGLLHFQVNGGDTPALAFVSFSSPSPGLQILDFALF 179
RiABP19.9       AGFISSDNTVYLKTLKKGDVMVFPQGLLHFQVNGGGTSALAFVSFSSPSPGLQILDFALF 178
RiABP19.1       SGFIDSNNKVYLKTLNKGDIIVIPPGLLHFQVNGGDTPVLEFAFFSSADPGVQILENSLF 179
RiABP19.2       AGFIDSNNKVYLKTLKQGDTMILPQGLLHFQKNGGDTPALLFAAFNSENPGVQLLEFALF 179
RiABP19.4       GGFIDSNNKVYLKTLKKGDIMIFPQGLLHFAKNEGDTPALLFAAFNSENPGVQFLENALF 179
RiABP19.5       AGFIDSNNKVYLKTLKKGDIMILPQGLLHFQKNGGDTPALVFTAYNSENPGVQLLETALF 179
RiABP19.6       AGFIDSNNKVYLKTLKKGDIMILPQGLLHFQKNGGDTPALLFAAFNSENPGVQLLENALF 178
RiABP19.3       GGFISSENRVYLKTLEKGDIMVFPQGLLHFQVNGGDTPALEFVSFSSDSPGLQILPNALF 79
RiABP19.7       AGFISSENKVYLKTLKKGDIMVFPRGLLHFQVNGGDTPALEFVSFSSDSPGLQLLPNALF 179
                .**: *:* ***:**.:** :::* **:**  * *.* .* *.  .* .**::.*  :**

RiABP19         KTDLSTELIAKTTLLDKSQIKKLKRLLGGTN--- 211
RiABP19.8       KNNLPTALIAQTTFLDVAQIKKLKGVLGGTN--- 210
RiABP19.9       KNNLPTSLVAATTFLDVAQIKKLKGVLGGTN--- 209
RiABP19.1       LSNLPTELIAQTTFLDTAQIKKLKDFLGGTN--- 210
RiABP19.2       QSDFPTELIAQTTLLDTAQIKKLKGLLGGTN--- 210
RiABP19.4       QSDFPTELIAQTTLLDTAQIKKLKGLLGGTN--- 210
RiABP19.5       QSDFPTELIAQTTLLDTAQIKKLKGLLGGTN--- 210
RiABP19.6       QSDFPTELIAQTTLLDTAQIKKLKGLLGGTN--- 209
RiABP19.3       LNNLPAELIAQSTFLDTAEIDQETQ---GSSWWY 110
RiABP19.7       QNNLPTELIAQSTFLDTAEIKRLKDLLGGTN--- 210
                 .:: : *:* :*:** ::*.: .    *:.
```

**Supplementary Figure S2:** Protein sequence alignment of RiABP19 and homologues located on chromosome 1 of red raspberry. Conserved GLP motifs are highlighted in blue boxes. The first box is important for formation of a disulphide bond and plays a role in protein-protein interactions. The second and third boxes are involved in metal ion binding [24]. Consensus is indicated below sequences where '*' indicates identical residues, ':' indicates amino acids with similar characteristics and '.' indicates amino acids with similar physiochemical properties.

```
G.Moy       GCAACTAAACACAAAACAAAATAAAAAAAACAACTTCATGAAATGAAGCTAGGCAACCTC    1141
Latham      --------ACACAAAACAAAAT----AAAAAAACTTCATGAAATGAAGCTAGGCAACCTC    1153
Consensus           **************    ****.******************************
G.Moy       AAAACTAGCCAAAATTAAAAAATTGATTAGTCTATTAGATGCTCTAAATCTTGCTCGTCC    1081
Latham      AAAACTAGCCAAAATTAAAAAATTGATTAGTCTATTAGATGCTCTAAATCTTGCTCGTCC    1093
Consensus   ************************************************************
G.Moy       AACATGGAAACTGAAAAGTAAAGAGCTTGATGAACAAGTAAAAGGCGTGTTAACCCACTA    1021
Latham      AACATGGAAACTGAAAAGTAAAGAGCTTGATGAACAAGTAAAAGGCGTGTTAACCCACTA    1033
Consensus   ************************************************************
G.Moy       ATCCGTCCACGTACTTGAACTCTCATCATATATCTGCATCTTGTAATTGGGGCCTTTCAT    961
Latham      ATCCGTCCACGTACGTGAACTCTCATCATATATCTGCATCTTGTAATTGGGGCCTTTCAT    973
Consensus   ************** *********************************************
G.Moy       GAATTAATTAGTATGCAAAGTATAGGACTAAACGCAGAGGATGGTGCTTCATGAATTGAG    901
Latham      GAATTAATTAGTATGCAAAGTATAGGACTAAACACAGAGGATGGTGCTTCATGAATTGAG    913
Consensus   *********************************.**************************
G.Moy       CTAGATCAGGGAGCAGAAAACCAATGGAAGAGAAGAGATCTAGAAAGCCATTATAATTGG    841
Latham      CTAGATCAGGGAGCAGAAAACCAATGGAAGAGAAGAGATCTAGAAAGCCATTATAATTGA    853
Consensus   ***********************************************************.
             b-2
G.Moy       AAAAAAAAGTGAGGGCTGGCCGGCTGGCCGGTCACCTCACATGTTTGTCAATAATTTCC     781
Latham      A-AAAAAAGTCAGGGCTGGCCGGCTGGTCGGTCACCTCACATGTTTGTCAATAATTTCC     794
Consensus   * ********* **************** ****************************** 
                                      a-5     a-4
G.Moy       GGCCTCGATCTGCTGTTACCTTGACTAGTTGACGTAATATATTTAAGCAAATGGCTGTAC    721
Latham      GGCCTCGATCTGCTGTTACCTTGACTAGTTGACGTAATATATTTAAGCAAATGGCTGTAC    734
Consensus   ************************************************************
G.Moy       GTATCGGTGGAAGTAGTCGCTGCAAGATCATAGATTAATTAACATGGAAACTTGGCTACA    661
Latham      GTATCGGTGGAAGTAGTCGCTGCAAGATCAT-------TTAACATGGAAACTTGGCTACA    681
Consensus   *******************************       **********************
G.Moy       CGTACAGC----TAGCGCGGGAATTAGTATTTCATAGACTAATTATGAGTTGCTTCCATA    605
Latham      CGTACAGCGCGCTAGCGCGGGAATTAGTATTTCATAGATTAATTATGAGTTGCTTCTATA    621
Consensus   ********    ************************* ***************** ***
                                                                  c-3
G.Moy       ATACGAATTCCTGTTAAATGATTATATAACTGGTTTGAAGCAATTTCGGGTCAAGTTTGT    545
Latham      ATACGAATTCCTGTTAAATGATTATACAACTGGTTTGAAGCAATTTCGGGTCAAGTTTGT    561
Consensus   ************************** *********************************
                        b-1         a-3
G.Moy       CTCTGGAAATTGATAAACTTAGTTGACGTGCGGTACGTGATCGAATTTGTCCCTTTCAAA    485
Latham      CTCTGGAAATTGAAAAACTTAGTTGACGTGCGGTACGTGATCGAATTTGTCCCTTTCAAA    501
Consensus   *************:**********************************************
G.Moy       AGTCTCCCCACTGTTTTAGATTTTATACTTTTACTCGTTGAGGACAATTTCTTGTTAATC    425
Latham      AGTCTCCCCGCTGTTTTAGATTTTATACTTTTACTCGTTGAGGATAATTTCTTGTTAATC    441
Consensus   *********.********************************** ***************
G.Moy       TGGCTAACTAGAGTTGATGTGTGAAATTTCTTTTTCTTCTTGAATTGGAGAAGATCTTGA    365
Latham      TGGCTAACTAGAGTTGATGTGTGAAATTTCTTTTTCTTCTTGAATTGAAGAAGATCTTGA    381
Consensus   ***********************************************.************
                                              c-2
G.Moy       TACGGCTTCCCCCCACCCCAATTTCCGTGCCTGTCTCGAAGACTACTTAATCCACAATTG    305
Latham      TACGGCTGCCCCCCACCCCAATTTCCGTGCCTGTCTCGAAGACTACTTAATCCACAATTG    321
Consensus   ******* ****************************************************
G.Moy       GATGGATTCAATGAACTAAATTAGCTAGTTTTAGAAACTTGTATATGGAATTCTTGTTGC    245
Latham      GATGGATTCAATGAACTAAATTAGCTAGTTCCAGAAACTTGTATATTGAATTCTTGTTGC    261
Consensus   ******************************  ************** *************
G.Moy       TTGTCTTTGAAGTCGTCGAAGTAGTAAGTAATCTGTTTCTCTCTCTTCTTATTGAATTCT    185
Latham      TTGTCTTTGAAGTCGTCGAAGTAGTAAGTAATCTGTTTCTCGATCTTCTTATTGAATTCT    201
Consensus   *****************************************  .*****************
                           a-2
G.Moy       TAGTCCTGCATGTTGACATCTTCTCACAACTAGAAGCTTCTTCCGTATACGTTTTATATA    125
Latham      TAGTCCT----GTTGACATCTTCTCACAACTAGAAGCTTCTTCCGTATACGTTTTATATC    145
Consensus   *******    ************************************************.
                                                a-1           d
G.Moy       -----------------------TATATATTATTTGACAGATATGAAGGCTGCCCCTCT     89
Latham      TATATATATATATATATATATATATATATATTATTTGACAGATATGAAGGCCGCCCCTCT    85
Consensus                          *************************** ********
                          c-1                  TATA box
G.Moy       TGTCAATTATTATTGTCTCAATCAATACGATTCAAGGTTATAAATAGCATTCAACAACTT    29
Latham      TGTCAATTATTATTGTCTCAATCAATACGATTCAAAGTTATAAATAGCATTCACCAACTT    25
Consensus   ***********************************.*****************.******
                      TSS
G.Moy       AGACCTTTACGTGCATATATATCATATC    1
Latham      AGACCTTTACGTGCATATCA----TATC    1
Consensus   ******************.:    ****
```

**Supplementary Figure S3**: Sequence of 1200 bp promoter region of *RiABP19* in Latham and G. Moy. Putative regulatory elements are either, underlined or highlighted. Auxin-responsive elements are highlighted in turquoise, and pathogenesis-related elements are indicated by letters and numbers (see Table 2), **a:** W-box, **b:** GT1 box, **c:** silencing element binding factor and **d:** GCC core. **TSS:** transcription starting site (red G). The putative TATA box is highlighted in yellow.

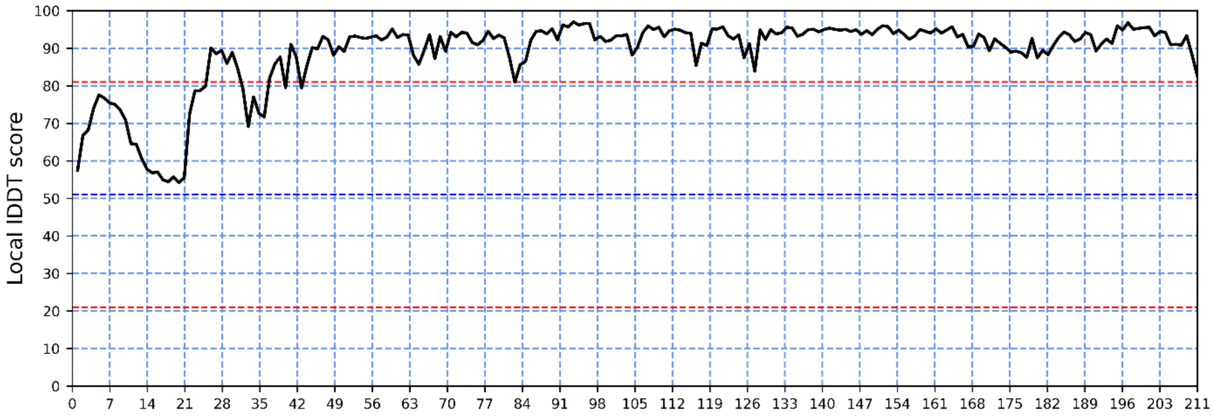


**Supplementary Figure S4:** Result of model quality assessment of RiABP19 monomer using the DeepUMQA method. The global local distance difference test (lDDT) score is 88.67 which is a score that assesses differences in local distances of all atoms within a protein structure. A score of lDDT above 80 is considered an accurate model.

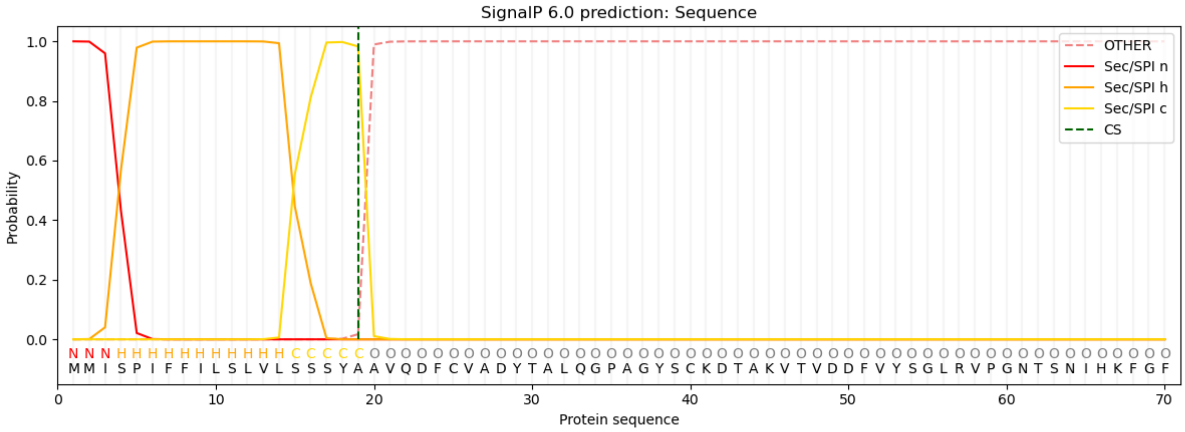


**Supplementary Figure S5:** SignalP-6.0 prediction of RiABP19. A signal peptide of 19 aa was predicted with a probability of 0.99. Location of peptide cleavage is predicted between 19 and 20 aa.

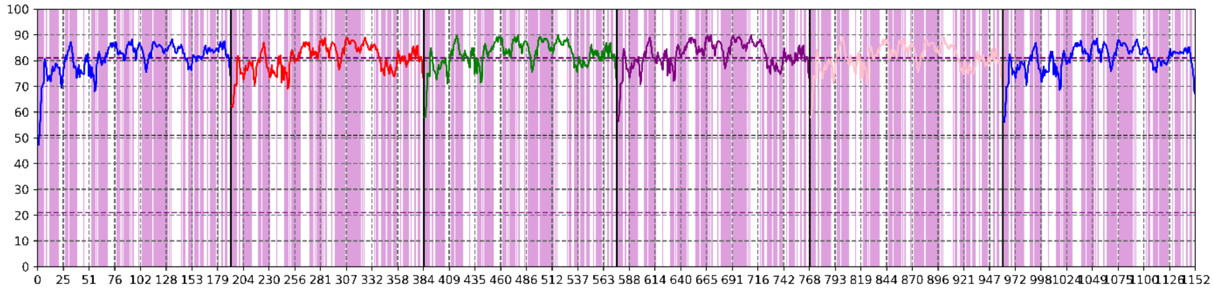


**Supplementary Figure S6:** Model quality assessment of RiABP19 hexamer using the DeepUMQA model. Per-residue lDDT is shown in the graph. The global lDDT score is 81.38. A score of above 80 is considered to represent an accurate model. Interface residues of each chain are provided in **Supplementary Table 3**.

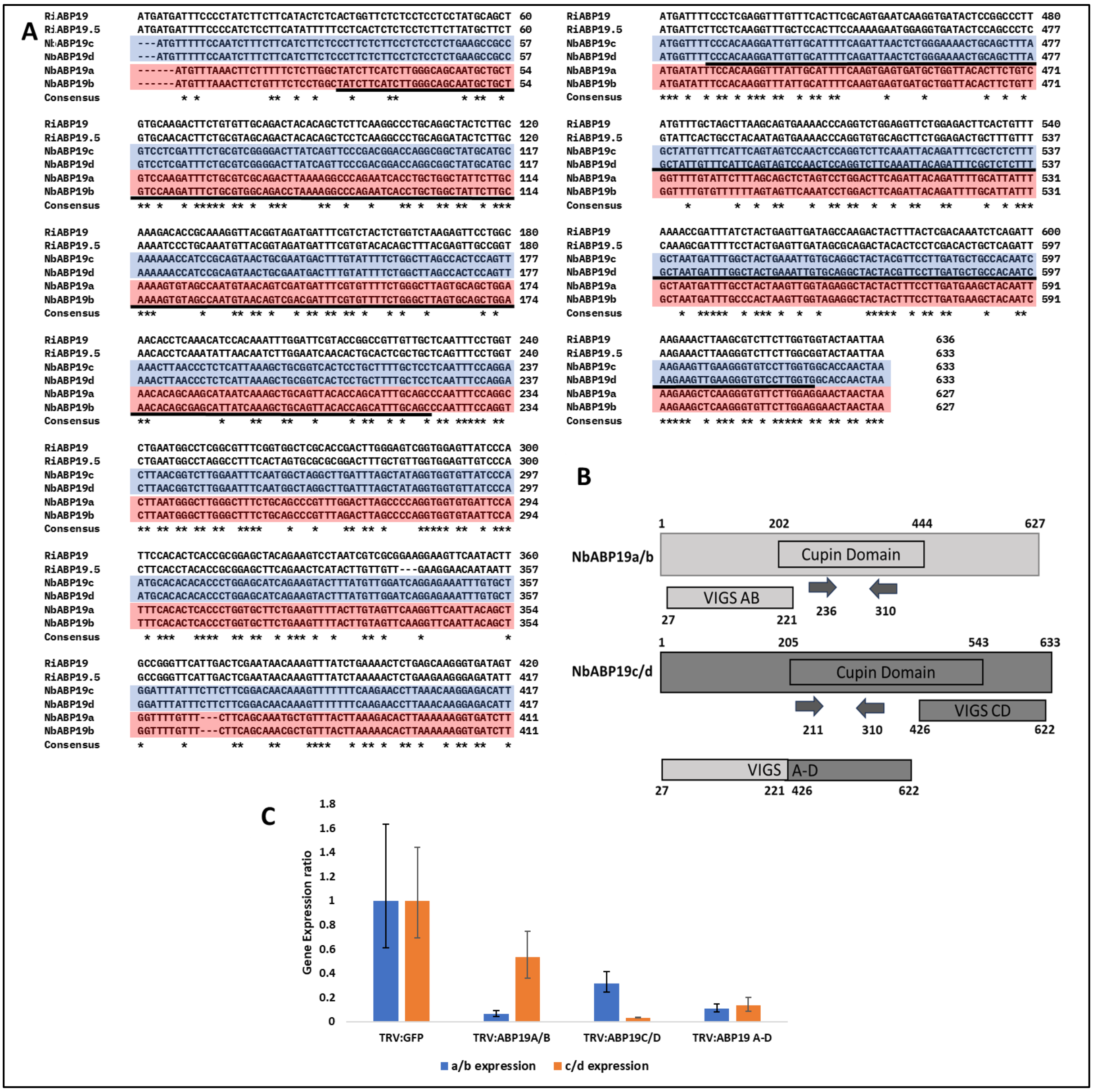


**Supplementary Figure S7: Virus-induced gene silencing of *NbABP19s* in *Nicotiana benthamiana* plants. A.** Nucleotide sequences of *RiABP19*, *RiABP19.5* and *NbABP19s*. The similarity in *NbABP19c/d* is highlighted in blue and the similarity in *NbABP19a/b* is highlighted in red. Silenced regions are indicated by black underlines. **B.** Schematic representation of VIGS constructs. The cupin domain and nucleotide regions selected for silencing are shown. **C.** Expression of *NbABP19A/B* and *NbABP19C/D* in silenced plants were compared to control *TRV:GFP*. Error bars are standard errors. Data was collected from two biological replicates.

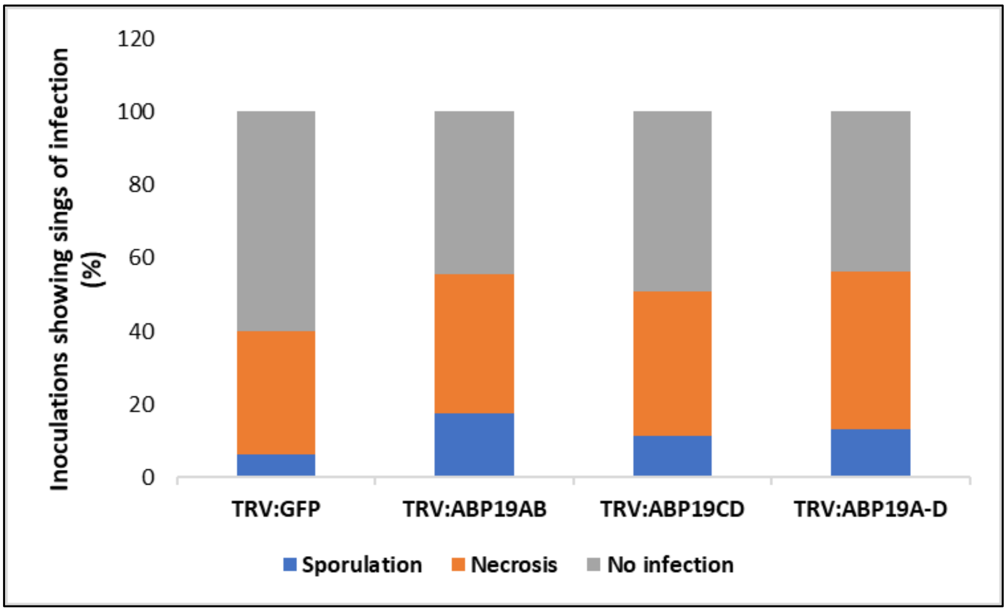

**Supplementary Figure S8:** Silencing of *NbABP19* results in increased susceptibility to *Phytophthora infestans* infection. Graph showing percentage of different signs of infection developing from *P. infestans* inoculations on *Nicotiana benthamiana* plants expressing different VIGS constructs at 12dpi. Data represents the combination of three independent experiments where the number of inoculations, n = >50.

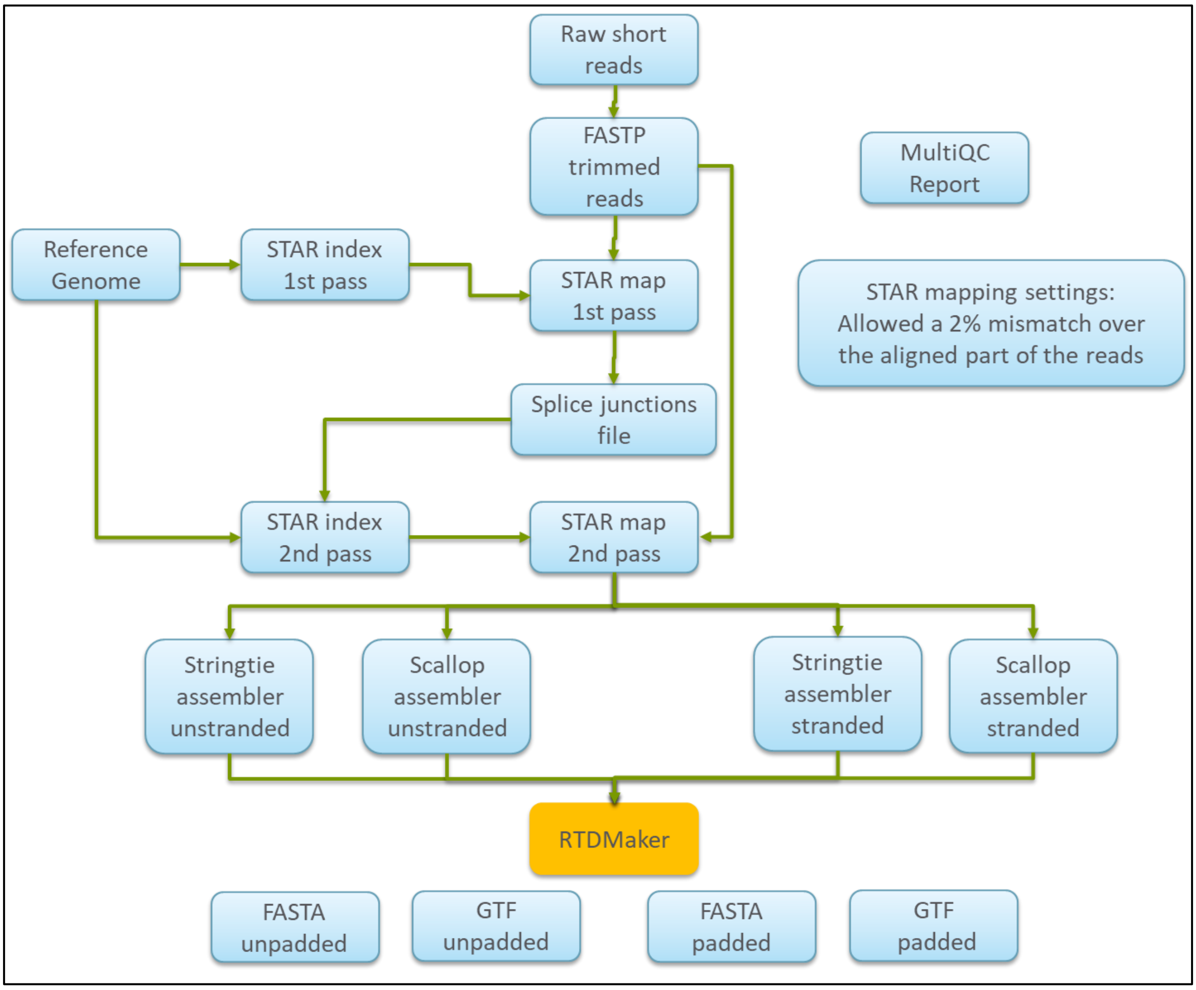


**Supplementary Figure S9:** Workflow of the pipeline used to develop reference transcript dataset (RTD). Illumina short reads (paired-ends) were used to run through the pipeline. Sequence reads were split at splicing sites and mapped to the reference genome.

**Supplementary Table S1**

List of differentially expressed genes (DEGs) categorised under 'defence response' in challenged Latham roots at 7 dpi. 1 $Log_2FC$= 2-fold change in expression.

| Gene ID | Annotation | $Log_2FC$ |
|---|---|---|
| rirtd4_HiC_scaffold_5G014910 | Protein P21-like | 4.8 |
| rirtd4_HiC_scaffold_7G030480 | Protein P21-like | 4.6 |
| rirtd4_HiC_scaffold_6G016490 | Thaumatin-like protein | 4.4 |
| rirtd4_HiC_scaffold_4G015780 | Chitinase 4-like | 4.1 |
| rirtd4_HiC_scaffold_367G000060 | Chitinase 4-like | 4 |
| rirtd4_HiC_scaffold_6G016560 | Thaumatin-like protein | 3.9 |
| rirtd4_HiC_scaffold_1G009040 | Pathogenesis related protein 3 | 3.5 |
| rirtd4_HiC_scaffold_2G014280 | Endochitinase-like | 2.9 |
| rirtd4_HiC_scaffold_2G002450 | Pathogenesis-related protein 1-like | 2.8 |
| rirtd4_ HiC_scaffold _3G031550 | Non-specific lipid-transfer protein | 2.5 |
| rirtd4_HiC_scaffold_2G030010 | Endochitinase 2-like | 1.4 |
| rirtd4_HiC_scaffold_6G018470 | Protein POLYCHOME-like | 1.3 |
| rirtd4_HiC_scaffold_2G038760 | Transport Inhibitor Response 1-like | 1.3 |
| rirtd4_HiC_scaffold_7G039220 | BON1-associated protein 2-like | 1.1 |
| rirtd4_HiC_scaffold_5G014910 | Thaumatin-like protein 1 | 1 |
| rirtd4_HiC_scaffold_4G027410 | Mitogen-activated protein kinase kinase NPK1 | 1 |
| rirtd4_HiC_scaffold_3G030040 | NB-ARC domain-containing disease resistance protein | -1 |
| rirtd4_HiC_scaffold_3G027250 | NB-ARC domain-containing disease resistance protein | -1 |
| rirtd4_HiC_scaffold_2G018100 | NB-ARC domain-containing disease resistance protein | -1 |
| rirtd4_HiC_scaffold_6G038400 | Disease resistance protein RGA3 | -1.1 |
| rirtd4_HiC_scaffold_2G031360 | Disease resistance protein (TIR-NBS-LRR class) | -1.1 |
| rirtd4_HiC_scaffold_6G026560 | NB-ARC domain-containing disease resistance protein | -1.1 |
| rirtd4_HiC_scaffold_3G053470 | NB-ARC domain-containing disease resistance protein | -1.1 |
| rirtd4_HiC_scaffold_3G026370 | NB-ARC domain-containing disease resistance protein | -1.1 |
| rirtd4_HiC_scaffold_5G042730 | NB-ARC domain-containing disease resistance protein | -1.1 |
| rirtd4_HiC_scaffold_2G025130 | Disease resistance protein (TIR-NBS-LRR class) | -1.1 |
| rirtd4_HiC_scaffold_2G052770 | Disease resistance protein At4g33300 | -1.1 |
| rirtd4_HiC_scaffold_2G000500 | Disease resistance protein (TIR-NBS-LRR class) | -1.1 |
| rirtd4_HiC_scaffold_2G024350 | Disease resistance protein (TIR-NBS-LRR class) | -1.2 |
| rirtd4_HiC_scaffold_6G041570 | NB-ARC domain-containing disease resistance protein | -1.2 |
| rirtd4_HiC_scaffold_3G022290 | NB-ARC domain-containing disease resistance protein | -1.2 |
| rirtd4_HiC_scaffold_3G016660 | Disease resistance protein (TIR-NBS-LRR class) | -1.2 |
| rirtd4_HiC_scaffold_7G017500 | NB-ARC domain-containing disease resistance protein | -1.2 |
| rirtd4_HiC_scaffold_7G035090 | Disease resistance protein (TIR-NBS-LRR class) | -1.2 |
| rirtd4_HiC_scaffold_6G038350 | Disease resistance protein RGA3 | -1.3 |
| rirtd4_HiC_scaffold_7G015950 | NB-ARC domain-containing disease resistance protein | -1.3 |
| rirtd4_HiC_scaffold_3G004650 | VQ motif containing protein | -1.3 |
| rirtd4_HiC_scaffold_6G050080 | Disease resistance protein (TIR-NBS-LRR class) | -1.3 |
| rirtd4_HiC_scaffold_5G001220 | Uncharacterized protein | -1.3 |
| rirtd4_HiC_scaffold_7G015950 | NB-ARC domain-containing disease resistance protein | -1.4 |
| rirtd4_HiC_scaffold_6G058690 | WRKY transcription factor 72 | -1.4 |

| | | |
|---|---|---|
| rirtd4_HiC_scaffold_2G040080 | Pathogenesis-related protein 5 | -1.4 |
| rirtd4_HiC_scaffold_4G044160 | MACPF domain-containing protein At1g14780 | -1.4 |
| rirtd4_HiC_scaffold_6G014080 | NB-ARC domain-containing disease resistance protein | -1.5 |
| rirtd4_HiC_scaffold_6G038400 | Disease resistance protein RGA3 | -1.6 |
| rirtd4_HiC_scaffold_3G030040 | NB-ARC domain-containing disease resistance protein | -1.6 |
| rirtd4_HiC_scaffold_3G021980 | NB-ARC domain-containing disease resistance protein | -1.7 |
| rirtd4_HiC_scaffold_6G014550 | MLO-like protein 6 | -1.7 |
| rirtd4_HiC_scaffold_1G022090 | Disease resistance protein (TIR-NBS-LRR class) | -1.7 |
| rirtd4_HiC_scaffold_7G016530 | NB-ARC domain-containing disease resistance protein | -1.8 |
| rirtd4_HiC_scaffold_7G033980 | NB-ARC domain-containing disease resistance protein | -1.8 |
| rirtd4_HiC_scaffold_6G014550 | MLO-like protein 6 | -1.8 |
| rirtd4_HiC_scaffold_3G019810 | NB-ARC domain-containing disease resistance protein | -1.9 |
| rirtd4_HiC_scaffold_1G008230 | Glucan endo-1,3-beta-glucosidase 6 | -2.1 |

### **Supplementary Table S2**

Characterised GLPs with accession numbers that were used to run sequence alignment and phylogenetic tree analysis.

| **Species** | **Abbreviation** | **Accession Number** |
|---|---|---|
| *Arachis hypogaea* | AhGLP5 | ADD71879.1 |
| *Arabidopsis thaliana* | AtGLP3a | P94072 |
| *Arabidopsis thaliana* | AtGLP5 | AAB51569 |
| *Beta vulgaris* | BvGLP1 | AAO85278 |
| *Gossypium hirsutum* | GhABP19 | QBF58681 |
| *Gossypium hirsutum* | GhGLP2 | QBF58682 |
| *Glycine max* | GmGLP10 | EU916258 |
| *Helianthus annuus* | HaGLP1 | AKM12467 |
| *Hordeum vulgare* | HvGER1a | ABG46232 |
| *Hordeum vulgare* | HvGER2a | ABG46233 |
| *Hordeum vulgare* | HvGER4d | ABG46236 |
| *Hordeum vulgare* | HvGER5a | ABG46237 |
| *Lolium perenne* | LpOXO1 | CAC19429 |
| *Oryza sativa* | OsGLP1 | AAC04836 |
| *Oryza sativa* | OsGLP3-7 | PRJNA808176 |
| *Pharbitis nil* | PnGLP | P45853 |
| *Prunus persica* | PpABP19 | U79114 |
| *Prunus persica* | PpABP20 | U81162 |
| *Prunus salicina* | PsABP2 | EU310511 |
| *Pisum sativum* | PsGER1 | CAB65369 |
| *Prunus salicina* | PsGLP1 | EU310513 |
| *Prunus salicina* | PsGLP2 | ACA03784 |
| *Rhododendron mucronatum* | RmGLP2 | BAG75123 |
| *Triticum aestivum* | TaGER2 | P15290 |
| *Triticum aestivum* | TaGER3 | P26759 |
| *Triticum aestivum* | TaGLP2a | CAB55558 |

| | | |
|---|---|---|
| *Vitis vinifera* | VvGLP3 | AAQ63185 |
| *Zea mays* | ZmGLP1 | AAQ95582 |

### Supplementary Table S3

Interface residue lDDT of homohexamer model of RiABP19.

| | | |
|---|---|---|
| Chain | A | A1:47.5 A2:51.97 A3:59.88 A4:69.17 A5:70.38 A6:71.02 A9:76.19 A12:74.09 A13:71.93 A14:74.93 A15:77.67 A16:75.98 A17:76.25 A18:78.82 A19:78.6 A20:77.54 A21:77.08 A28:82.0 A29:80.66 A30:83.55 A32:85.71 A33:87.22 A34:85.18 A35:81.91 A36:77.38 A37:77.81 A38:77.58 A39:73.22 A46:74.67 A54:77.35 A55:73.95 A56:74.22 A57:68.04 A59:78.72 A61:83.47 A62:81.98 A63:79.39 A64:78.24 A65:80.05 A66:80.21 A67:79.15 A68:82.92 A69:81.34 A79:84.48 A80:82.29 A81:82.74 A82:83.87 A84:82.84 A86:80.07 A87:80.12 A88:83.39 A89:85.75 A90:86.25 A91:87.3 A93:86.54 A95:84.05 A101:86.8 A104:84.35 A105:82.8 A106:81.08 A107:78.3 A108:75.58 A109:77.68 A110:80.22 A111:82.06 A112:82.98 A113:80.96 A114:85.27 A116:87.32 A117:86.6 A118:85.53 A119:85.26 A120:85.88 A121:86.06 A122:85.76 A123:85.54 A124:84.99 A125:83.31 A126:82.79 A127:84.4 A128:83.91 A129:83.62 A131:85.78 A133:87.53 A143:86.09 A145:82.71 A146:82.84 A147:80.73 A150:76.59 A151:77.7 A152:77.73 A153:76.26 A154:79.89 A156:81.31 A157:81.22 A158:79.62 A159:81.77 A160:81.78 A162:81.27 A163:80.42 A164:81.55 A165:81.96 A169:82.7 A172:83.53 A173:84.21 A174:84.27 A175:83.1 A176:83.33 A177:84.0 A180:83.4 A183:85.58 A184:84.25 A187:83.14 A188:81.17 |
| Chain | B | B1:62.46 B2:61.89 B3:64.21 B4:70.42 B5:70.68 B6:70.48 B9:78.95 B12:74.94 B13:72.66 B14:75.05 B15:78.04 B16:76.28 B17:76.31 B18:79.03 B19:79.73 B20:78.49 B21:78.05 B28:80.79 B29:79.41 B30:82.72 B32:84.75 B33:86.97 B34:84.91 B35:81.12 B36:77.51 B37:77.05 B38:75.96 B39:73.41 B46:73.48 B54:77.27 B55:74.73 B56:75.16 B57:69.13 B59:79.52 B61:84.03 B62:83.19 B63:80.83 B64:79.35 B65:80.69 B66:80.78 B67:80.03 B68:83.13 B69:81.71 B79:83.81 B80:80.85 B81:82.18 B82:84.87 B84:82.88 B86:81.07 B87:80.82 B88:84.21 B89:87.48 B90:87.17 B91:88.16 B93:87.18 B95:84.58 B101:87.08 B104:85.64 B105:83.93 B106:82.58 B107:79.4 B108:76.44 B109:77.86 B110:80.36 B111:82.91 B112:84.75 B113:82.46 B114:86.69 B116:88.9 B117:88.11 B118:86.48 B119:86.35 B120:87.67 B121:87.86 B122:87.92 B123:87.44 B124:86.98 B125:84.56 B126:83.47 B127:85.05 B128:84.23 B129:82.97 B131:86.36 B133:88.13 B143:86.34 B145:84.17 B146:84.35 B147:82.69 B150:75.86 B151:76.81 B152:76.21 B153:74.24 B154:79.29 B156:77.96 B157:76.23 B158:74.81 B159:78.29 B160:77.06 B162:74.24 B163:73.78 B164:75.81 B165:77.46 B169:78.94 B172:79.77 B173:81.7 B174:79.53 B175:77.58 B176:79.48 B177:79.19 B180:80.08 B183:82.35 B184:80.64 B187:80.03 B188:79.61 |
| Chain | C | C1:57.95 C2:58.16 C3:65.1 C4:74.88 C5:74.81 C6:73.77 C9:80.56 C12:75.06 C13:72.63 C14:75.2 C15:78.42 C16:76.51 C17:77.07 C18:79.48 C19:80.13 C20:79.05 C21:79.32 C28:84.27 C29:82.12 C30:86.05 C32:88.34 C33:89.61 C34:88.4 C35:85.61 C36:82.39 C37:82.4 C38:82.14 C39:80.31 C46:77.2 C54:80.63 C55:78.1 C56:77.77 C57:71.44 C59:81.89 C61:86.55 C62:85.51 C63:82.86 C64:81.35 C65:83.19 C66:83.71 C67:83.57 C68:85.34 C69:84.39 C79:85.11 C80:81.96 C81:82.44 C82:84.54 C84:83.2 C86:81.53 C87:81.16 C88:85.05 C89:88.19 C90:87.92 C91:89.18 C93:88.35 C95:85.86 C101:87.6 C104:85.76 C105:83.35 C106:82.18 C107:78.88 C108:75.67 C109:77.0 C110:79.63 C111:82.08 C112:83.92 C113:81.77 C114:87.0 C116:89.3 C117:88.61 C118:87.18 C119:87.15 C120:88.48 C121:88.0 C122:88.44 C123:87.97 C124:87.25 C125:84.54 C126:83.76 C127:85.28 C128:84.51 C129:82.86 C131:86.57 C133:88.61 C143:88.75 C145:85.23 C146:85.57 C147:84.19 C150:77.21 C151:78.21 C152:77.21 C153:75.43 C154:79.91 C156:78.88 C157:77.86 C158:77.21 C159:80.22 C160:79.41 C162:78.13 C163:77.32 C164:78.52 C165:80.18 C169:81.25 C172:81.9 C173:83.69 C174:82.51 C175:79.91 C176:81.37 C177:79.85 C180:80.5 C183:83.32 C184:81.21 C187:81.24 C188:80.69 |
| Chain | D | D1:58.26 D2:56.36 D3:61.1 D4:70.98 D5:71.03 D6:71.79 D9:80.68 D12:75.57 D13:73.32 D14:75.74 D15:78.96 D16:77.01 D17:77.38 D18:80.04 D19:80.51 D20:79.61 D21:79.43 D28:83.44 D29:80.94 D30:85.31 D32:87.38 D33:89.0 D34:87.3 D35:84.27 D36:80.93 D37:80.93 D38:80.8 D39:79.28 D46:76.42 D54:80.33 D55:78.2 D56:77.99 D57:72.1 D59:81.82 D61:86.06 D62:85.48 D63:82.89 D64:81.07 D65:82.63 D66:83.18 D67:82.76 D68:85.35 D69:84.15 D71:87.04 D79:85.03 D80:82.33 D81:83.2 D82:85.34 D84:83.76 |

| | | |
|---|---|---|
| | | D85:84.76 D86:82.04 D87:81.85 D88:85.65 D89:88.88 D90:88.58 D91:89.65 D93:88.56 D95:85.71 D101:87.56 D104:86.58 D105:84.38 D106:83.56 D107:80.1 D108:76.92 D109:78.31 D110:80.96 D111:82.95 D112:85.08 D113:82.46 D114:87.23 D116:89.49 D117:88.67 D118:87.11 D119:87.23 D120:88.85 D121:88.8 D122:89.18 D123:88.52 D124:88.04 D125:85.78 D126:84.69 D127:86.21 D128:85.78 D129:83.82 D131:86.9 D133:88.66 D143:88.23 D145:85.37 D146:85.82 D147:84.3 D150:77.0 D151:78.04 D152:76.93 D153:74.93 D154:79.67 D156:78.37 D157:76.93 D158:75.85 D159:78.91 D160:77.54 D162:75.55 D163:74.69 D164:76.13 D165:77.96 D169:79.49 D172:80.38 D173:82.19 D174:80.43 D175:78.19 D176:79.91 D177:78.89 D180:79.9 D183:82.45 D184:80.53 D187:80.23 D188:79.71 |
| Chain | E | E1:58.0 E2:58.72 E3:64.98 E4:74.0 E5:73.93 E6:73.0 E9:80.45 E12:75.55 E13:72.99 E14:75.57 E15:78.53 E16:76.66 E17:76.69 E18:79.62 E19:80.69 E20:78.93 E21:78.58 E28:83.75 E29:81.83 E30:85.52 E32:87.45 E33:89.17 E34:87.38 E35:84.57 E36:80.89 E37:80.81 E38:80.5 E39:78.24 E46:74.59 E54:79.12 E55:76.96 E56:77.16 E57:71.14 E59:81.37 E61:85.59 E62:84.69 E63:81.88 E64:80.32 E65:81.91 E66:82.22 E67:81.7 E68:84.3 E69:82.89 E79:84.14 E80:80.71 E81:81.62 E82:84.06 E84:83.05 E85:83.95 E86:82.05 E87:81.43 E88:84.69 E89:87.52 E90:87.2 E91:88.06 E93:87.09 E95:84.65 E101:86.67 E104:85.06 E105:82.89 E106:81.69 E107:78.69 E108:75.5 E109:76.73 E110:79.19 E111:81.53 E112:82.89 E113:80.67 E114:85.83 E116:88.32 E117:87.47 E118:85.96 E119:85.82 E120:86.72 E121:86.83 E122:86.89 E123:86.83 E124:86.33 E125:83.9 E126:83.36 E127:84.94 E128:83.54 E129:82.11 E131:85.61 E133:87.77 E143:86.3 E145:83.52 E146:83.8 E147:81.65 E150:75.82 E151:76.94 E152:76.6 E153:74.61 E154:79.5 E156:77.04 E157:75.16 E158:74.94 E159:78.64 E160:77.12 E162:74.75 E163:74.89 E164:76.09 E165:77.81 E169:79.0 E172:80.11 E173:82.08 E174:80.24 E175:77.95 E176:79.13 E177:78.23 E180:78.88 E183:81.23 E184:79.31 E187:79.41 E188:78.83 |
| Chain | F | F1:57.79 F2:56.16 F3:60.8 F4:69.9 F5:69.91 F6:70.93 F9:78.8 F12:75.69 F13:73.22 F14:75.78 F15:78.33 F16:76.41 F17:76.56 F18:79.5 F19:79.54 F20:78.75 F21:78.19 F28:82.1 F29:80.0 F30:83.45 F32:85.49 F33:86.89 F34:84.59 F35:80.95 F36:76.83 F37:76.69 F38:75.45 F39:71.71 F46:75.82 F54:77.51 F55:74.56 F56:74.76 F57:68.57 F59:79.18 F61:83.75 F62:82.4 F63:79.99 F64:78.51 F65:80.04 F66:80.23 F67:79.11 F68:82.9 F69:81.02 F79:85.33 F80:82.7 F81:83.27 F82:84.49 F84:82.69 F85:82.63 F86:79.55 F87:79.98 F88:83.49 F89:86.03 F90:86.53 F91:87.8 F93:86.92 F95:83.94 F101:87.3 F104:84.62 F105:83.0 F106:81.24 F107:78.63 F108:75.94 F109:77.29 F110:79.66 F111:82.11 F112:83.19 F113:81.91 F114:86.04 F116:87.99 F117:87.17 F118:85.85 F119:85.74 F120:86.72 F121:86.9 F122:86.61 F123:85.86 F124:85.14 F125:83.64 F126:82.56 F127:83.82 F128:83.74 F129:83.12 F131:85.68 F133:87.75 F143:85.31 F145:82.49 F146:82.34 F147:79.96 F150:76.36 F151:77.29 F152:77.31 F153:75.39 F154:78.45 F156:79.94 F157:79.8 F158:77.91 F159:79.88 F160:80.01 F161:79.04 F162:78.92 F163:78.41 F164:79.18 F165:80.22 F169:81.71 F172:82.88 F173:82.96 F174:83.12 F175:82.81 F176:82.74 F177:83.51 F180:82.76 F183:83.58 F184:82.83 F187:81.72 F188:79.59 |

### Supplementary Table S4

Public and in-house RNA-seq datasets used to assemble a raspberry reference transcript database.

| **Project ID** | **Cultivars** | **Total Gbp** | **Tissues** | **Conditions** |
|---|---|---|---|---|
| **PRJNA476755** | Joan J. | 131 | Fruit (ovary wall, seed/ovule) | 0 and 12 DPA |
| **PRJEB28528** | Glen Dee | 8 | Leaf | Virus infection |
| **PRJNA354231** | Amira | 46 | Leaf | ToSRV infected and non-infected |
| **PRJNA560307** | Wakefield, Heritage | 6 | Axillary buds | n/a |
| **PRJEB73351** | Glen Moy, Latham | 156 | Root tips | Root infected and non-infected |

| | | | | |
|---|---|---|---|---|
| **In-house** | 22 varieties | 134 | Whole plant | n/a |
| **In-house** | Glen Dee | 150 | Root | Root rot infected, non-infected, T0, T1, T10 |
| **In-house** | Glen Dee/Glen Ample | 631 | Buds | Dormancy |

### Supplementary Table S5

Primer sequences used for cloning and RT-qPCR.

| **Primer Name** | **Primer Sequence (5'-3')** | **Application** |
|---|---|---|
| RiABP19_F | CACTTCGCAGTGAATCAAGG | RT-qPCR |
| RiABP19_R | AGTCTCCAGAACCTCCAGAC | RT-qPCR |
| RiABP19.5_F | TCCCTAGCTAACGCGGACTT | RT-qPCR |
| RiABP19.5_R | CTGGTGTGTGTGTGTGCATTG | RT-qPCR |
| RiABP19.7_F | AACCTCTTTTCTTAGTGGTGGTTC | RT-qPCR |
| RiABP19.7_R | TTGATGGTAATATTTTTGACAACTAGC | RT-qPCR |
| RiEF1a_qRT_F | AGGAGCCCAAGTTCTTGAAGA | RT-qPCR |
| RiEF1a_qRT_R | CCTCACAGCAAAACGACCAA | RT-qPCR |
| Nb_qRT_ABP19A/B_F | TTAATGGGCTTGGGCTTTCT | RT-qPCR |
| Nb_qRT_ABP19A/B_R | CAGGGTGAGTGTGAAATGGA | RT-qPCR |
| Nb_qRT_ABP19CDE_F | CCTGCTTTTGCTCCTCAATTT | RT-qPCR |
| Nb_qRT_ABP19CDE_R | GGTGTGTGTGCATTGGGATA | RT-qPCR |
| GW_Latham_RiABP19.5_ F | AAA-GCA-GGC-TTC-ACC-ATG-ATGATTTTCCCCATCTCC | GW cloning |
| GW_Latham_RiABP19.5_ R | GAA-AGC-TGG-GTC-ATTAGTACCGCCAAGAAGACC | GW cloning |
| GW_Latham_RiABP19_ F | AAA-GCA-GGC-TTC-ACC-ATG-ATGATTTCCCCTATCTT | GW cloning |
| GW_Latham_RiABP19_ R | GAA-AGC-TGG-GTC-ATTAGTACCACCAAGAAGACG | GW cloning |
